# Aplp1 and the Aplp1-Lag3 Complex facilitates transmission of pathologic α-synuclein

**DOI:** 10.1101/2021.05.01.442157

**Authors:** Xiaobo Mao, Hao Gu, Donghoon Kim, Yasuyoshi Kimura, Ning Wang, Enquan Xu, Haibo Wang, Chan Chen, Shengnan Zhang, Chunyu Jia, Yuqing Liu, Hetao Bian, Senthilkumar S. Karuppagounder, Longgang Jia, Xiyu Ke, Michael Chang, Amanda Li, Jun Yang, Cyrus Rastegar, Manjari Sriparna, Preston Ge, Saurav Brahmachari, Sangjune Kim, Shu Zhang, Yasushi Shimoda, Martina Saar, Creg J. Workman, Dario A. A. Vignali, Ulrike C. Muller, Cong Liu, Han Seok Ko, Valina L. Dawson, Ted M. Dawson

## Abstract

Pathologic α-synuclein (α-syn) spreads from cell-to-cell, in part, through binding to the lymphocyte-activation gene 3 (Lag3). Here we report that amyloid β precursor-like protein 1 (Aplp1) forms a complex with Lag3 that facilitates the binding, internalization, transmission, and toxicity of pathologic α-syn. Deletion of both Aplp1 and Lag3 eliminates the loss of dopaminergic neurons and the accompanying behavioral deficits induced by α-syn preformed fibrils (PFF). Anti-Lag3 prevents the internalization of α-syn PFF by disrupting the interaction of Aplp1 and Lag3, and blocks the neurodegeneration induced by α-syn PFF *in vivo*. The identification of Aplp1 and the interplay with Lag3 for α-syn PFF induced pathology advances our understanding of the molecular mechanism of cell-to-cell transmission of pathologic α-syn and provides additional targets for therapeutic strategies aimed at preventing neurodegeneration in Parkinson’s disease and related α-synucleinopathies.

**One Sentence Summary:** Aplp1 forms a complex with Lag3 that facilitates the binding, internalization, transmission, and toxicity of pathologic α-synuclein.

**Graphical Abstract:** Aplp1 and the Aplp1-Lag3 complex facilitates transmission of pathologic α-synuclein. Aplp1 is a receptor that drives pathologic α-syn transmission, and genetic depletion of Aplp1 can significantly reduce the α-synuclein pathogenesis. Aplp1 and Lag3 forms an Aplp1-Lag3 complex that accounts for substantial binding of pathologic α-syn to cortical neurons. Together Aplp1 and Lag3 play a major role in pathologic α-syn internalization, transmission and toxicity. Double knockout of Aplp1 and Lag3 and or a Lag3 antibody that disrupts the Aplp1 and Lag3 complex almost completely blocks α-syn PFF-induced neurodegeneration.

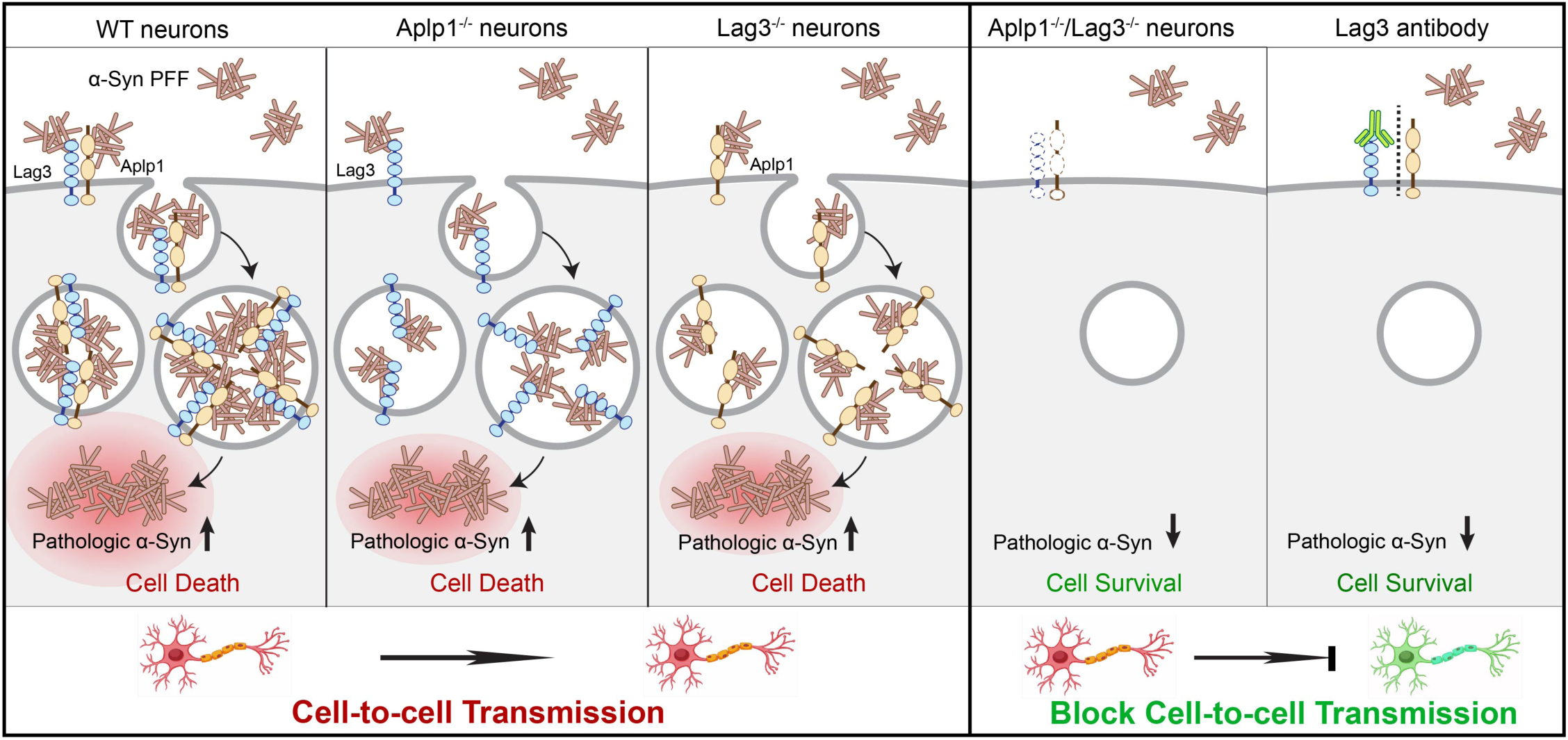

## INTRODUCTION

α-Synucleinopathies are a subset of neurodegenerative diseases (*1*), including Parkinson’s disease (PD), dementia with Lewy Bodies (DLB), and multiple system atrophy (MSA), which are characterized by abnormal accumulation of misfolded α-synuclein (α-syn) in neurons or glial cells. Pathologic α-syn, which is a prion-like protein, progressively spreads from the enteric and peripheral nervous systems into the central nervous system (*2*). Support for pathologic α-syn transmission is the appearance of Lewy body (LB) pathology in transplanted fetal mesencephalic dopaminergic neurons of PD patients (*3–5*). In cell-to-cell transmission models of pathologic α-syn, exogenous recombinant α-syn preformed fibrils (PFF) enter neurons and seed endogenous α-syn monomers (*6*), leading to substantial α-syn pathology and neurotoxicity (*7*). Intrastriatal or gut injection of α-syn PFF induces pathologic α-syn spreading, resulting in PD-like motor and non-motor symptoms (*8–10*).

Lymphocyte-activation gene 3 (Lag3) is a receptor that facilitates α-syn transmission from neuron-to-neuron (*11*). Depletion of Lag3 significantly reduces the internalization of α-syn PFF in neurons, but fails to completely prevent the internalization of α-syn PFF, suggesting that other receptors and mechanisms for α-syn transmission likely exist (*11–17*). Although amyloid β precursor-like protein 1 (Aplp1) is a pathologic α-syn binding protein (*11*) that belongs to the conserved amyloid precursor protein (App) family, and is associated with neurodegenerative disorders (*18–21*), the role of Aplp1 in pathologic α-syn transmission and pathogenesis is not known. Here we show that genetic deletion of Aplp1 significantly inhibit the internalization of pathologic α-syn, cell-to-cell transmission, neurotoxicity and behavioral deficits induced by α-syn PFF. Furthermore, Aplp1 and Lag3 form a complex that drives pathologic α-syn transmission. Deletion of both Aplp1 and Lag3 eliminates the loss of dopaminergic (DA) neurons and the accompanying behavioral deficits induced by α-syn PFF. Disruption of the Aplp1/Lag3 complex by using an anti-Lag3 antibody prevents pathologic α-syn transmission and pathogenesis providing a novel route to prevent the degenerative process set in motion by pathologic α-syn. These findings indicate that Aplp1 plays a role in the cell-to-cell transmission of pathologic α-syn and provides a proof of concept for Lag3-targeting immunotherapy. These observations may lead to optimization of pathologic α-syn receptor targeted therapies.

## RESULTS

### α-Syn PFF binds to amyloid β precursor-like protein 1 (Aplp1)

We conjugated the α-syn monomer with biotin (α-syn-biotin) (*11*) and generated α-syn-biotin PFF using an established protocol (*10*). We confirmed that α-syn-biotin PFF specifically bind to Aplp1-transfected SH-SY5Y cells in a saturable manner, with a *K*_d_ of 430 nM (fig. S1A) by a well-established cell surface-binding assay as previously reported (*11*). In contrast, the α-syn-biotin monomer at concentration as high as 3,000 nM does not exhibit any appreciable binding to Aplp1-transfected SH-SY5Y cells (fig. S1A). β-Amyloid-biotin PFF binds to SH-SY5Y cells expressing Aplp1 at high concentrations in a non-specific manner, and β-amyloid-biotin monomer does not exhibit any appreciable binding signal (fig. S1B). Moreover, α-syn-biotin PFF does not bind to the amyloid precursor protein (App) or the amyloid precursor-like protein 2 (Aplp2)^11^, suggesting that the binding between Aplp1 and α-syn-biotin PFF is specific.

Like other App family proteins, Aplp1 is a single-pass transmembrane protein implicated in synaptic adhesion (*22*). Aplp1 possesses a large N-terminal extracellular domain and a small intracellular C-terminal region (*23*). To identify the α-syn PFF-binding domain, we sequentially deleted each of the four domains of Aplp1 (ΔE1, ΔAcD+E2, ΔJMR (juxtamembrane region), and ΔICD (intracellular domain)) (Fig. 1A). The membrane locations of different deletion mutants of FLAG-Aplp1 were determined first (fig. S1C and S1E) along with Lag3-Myc (fig. S1D and S1E) with the membrane marker wheat germ agglutinin (WGA) conjugated with Alexa Fluor^TM^ 488. This was followed by determining the binding of α-syn-biotin PFF to the Aplp1 deletion mutants by a cell surface-binding assay (Fig. 1B). These experiments suggest that: (i) α-syn-biotin PFF preferentially bind to the E1 domain, (ii) deletion of the AcD+E2 domain does not interfere with binding, (iii) deletion of JMR and the transmembrane (TM) domain substantially weakens the binding and serve as controls as these mutants lack cell membrane expression, and (iv) deletion of ICD mildly reduces the binding (Fig. 1B). Since α-syn-biotin PFF binding to Aplp1 is eliminated by deletion of the E1 domain, we further examined the two subdomain deletion mutants of ectodomain E1, the growth factor-like domain (ΔGFLD) and the copper-binding domain (ΔCuBD). Deletion of CuBD does not interfere with binding, while deletion of GFLD subdomain moderately interferes with the binding (Fig. 1B).

**Fig. 1.**
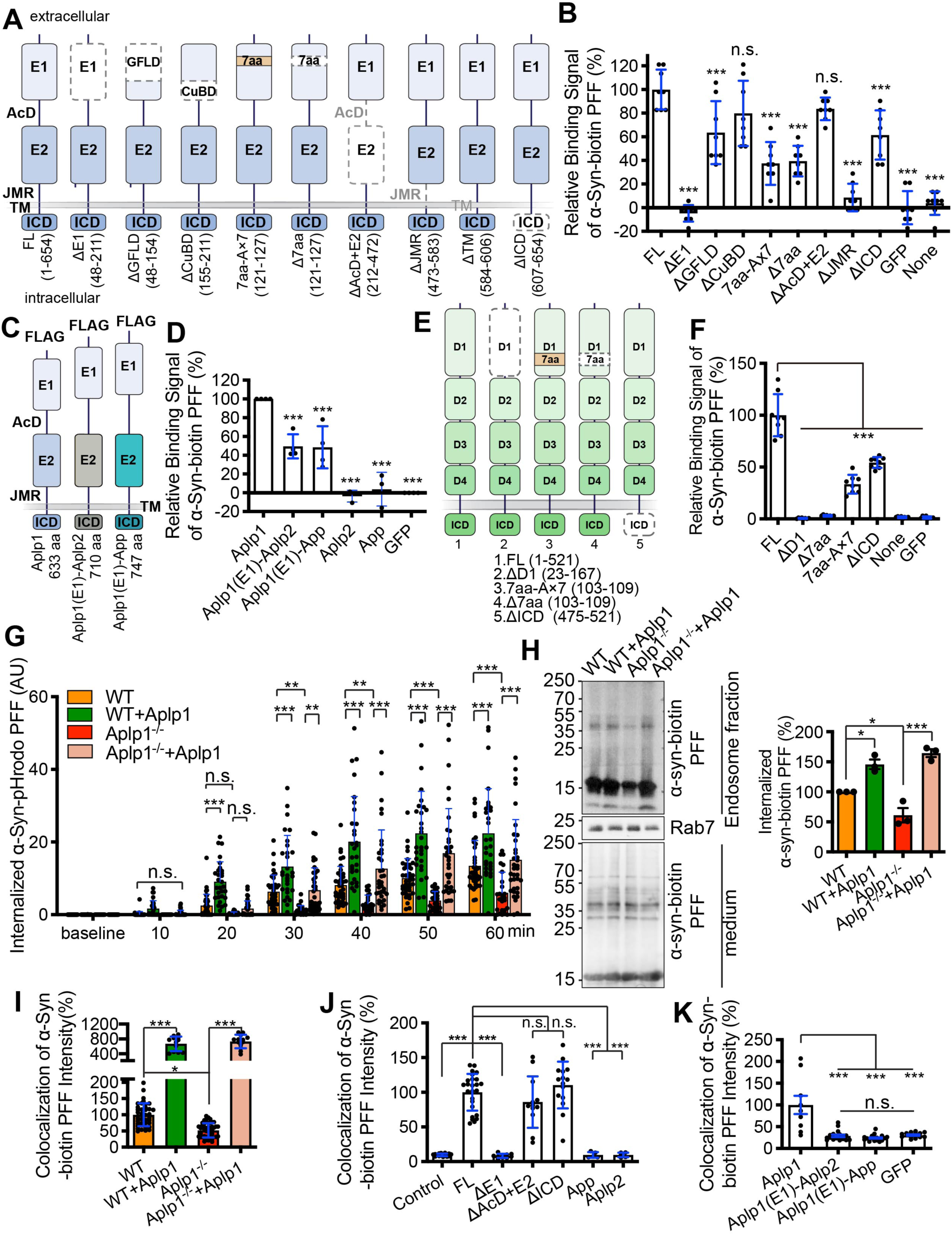
α-Syn PFF binds to Aplp1, and endocytosis of α-syn PFF is dependent on Aplp1. **(A)** Schematic diagram of Aplp1 deletions mutants: the ΔE1, ΔAcD+E2, ΔJMR (juxtamembrane region), ΔTM (transmembrane region), ΔICD (intracellular domain), two sub-domain deletion mutants of ectodomain E1 (ΔGFLD and ΔCuBD), and the deletion mutant (Δ7aa) and the substitutional mutant (7aa-A×7) of a seven-amino acid (7aa) motif. **(B)** Quantification of binding signals of deletion mutants of Aplp1 with α-syn-biotin PFF by normalization with the expression of indicated deletion mutants of Aplp1. **(C)** Schematic diagram of Aplp1 chimeras. **(D)** Quantification of binding signals of Aplp1(E1)-Aplp2 and Aplp1(E1)-App chimeras with α-syn-biotin PFF by normalization with the expression of indicated Aplp1 chimeras. Relative binding signals normalized to GFP (0%) and Aplp1 (100%) are shown. **(E)** Schematic diagram of Lag3 and deletions mutants. **(F)** Quantification of binding signals of deletion mutants of Lag3 with α-syn-biotin PFF by normalization with the expression of indicated deletion mutants of Lag3. **(B, D, F)** \*\*\**P* < 0.001, n.s., not significant. Data are the means ± SEM, from 3 individual experiments, one-way ANOVA followed by Dunnett’s correction. **(G)** Quantification of the intensity of the endocytosis of α-syn-pHrodo PFF in wildtype and Aplp1^−/−^ neurons by control lentivirus (WT: 38 cells, Aplp1^-/-^: 40 cells) and Aplp1-lentivirus transduction (WT+Aplp1: 29 cells, Aplp1^-/-^+Aplp1: 35 cells) with live image analysis from 3 independent experiments. Two way-ANOVA with Tukey’s correction. **(H)** Immunoblot (left panel) and quantification analysis (right panel) of α-syn-biotin PFF in the endosome fraction. Data are as means ± SEM. One way-ANOVA with Tukey’s correction. **(I)** Quantification of the intensity of the colocalization of α-syn-biotin PFF with Rab7 in WT and Aplp1^−/−^ neurons. WT (38 cells), WT+Aplp1 (12 cells), Aplp1^−/−^ (49 cells), Aplp1^−/−^+Aplp1 (13 cells) from 3 independent experiments. One way-ANOVA with Tukey’s correction. **(J)** Quantification of co-localization of α-syn-biotin PFF and Rab7 in WT neurons with transient expression of full-length Aplp1 (FL: 25 cells) and deletion mutants: ΔE1(10 cells), ΔAcD+E2(12 cells), ΔICD(15 cells), App(5 cells) and Aplp2(5 cells), and control (16 cells) from 3 independent experiments. One-way ANOVA followed by Dunnett’s correction. **(K)** Quantification of co-localization signal of α-syn-biotin PFF in WT neurons with transient expression of Aplp1, chimeric Aplp1(E1)-Aplp2 and Aplp1(E1)-App. One-way ANOVA followed by Tukey’s correction. **(G, I, J, K)** \**P* < 0.05, \*\**P* < 0.01, \*\*\**P* < 0.001, n.s., not significant, Data are means ± SEM.

To confirm that E1 domain of Aplp1 interacts with α-syn-biotin PFF, we constructed Aplp1 chimeras, in which E1-AcD of Aplp2 or App is replaced with the E1-AcD ectodomain of Aplp1 (Fig. 1C). Both Aplp1(E1)-Aplp2 and Aplp1(E1)-App exhibit binding to α-syn-biotin PFF, while Aplp2 and App do not bind to α-syn-biotin PFF (Fig. 1D). Since the binding of α-syn-biotin PFF to the Aplp1(E1)-Aplp2 and Aplp1(E1)-App chimeras does not bind at the same level as Aplp1 (Fig. 1D), the ICD subdomain of Aplp1 may help facilitate the interaction, which is consistent with effect of deletion of ICD subdomain of Aplp1 on α-syn-biotin PFF binding to Aplp1.

We identified a seven-amino acid (7aa) motif with a similar sequence in Aplp1 and Lag3: GGTRSGR (121–127) in Aplp1 and the GGLRSGR (103–109) in Lag3. Interestingly this 7aa motif is not in App and Aplp2, which might account for their lack of binding to pathologic α-syn. The 7aa motif is located in the α-syn-biotin PFF-binding domains of Aplp1 (GFLD subdomain) (Fig. 1A) and Lag3 (D1 domain) (Fig. 1E). Deletion of these 7 aa (Δ7aa) or the substitution with seven alanine (7aa-A×7) of Aplp1 or Lag3 significantly reduced the binding to both receptors (Fig. 1B and 1F). Taken together, these results suggest that the E1 domain of Aplp1 is the major binding domain for α-syn-biotin PFF, particularly in the GFLD subdomain and the 7aa motif in the E1 domain.

### Aplp1 mediates the endocytosis of α-syn PFF

To determine whether Aplp1 is involved in the endocytosis of α-syn PFF, we conjugated α-syn PFF to a pH-sensitive pHrodo red dye (α-syn-pHrodo PFF). As the pHrodo dye is non-fluorescent at neutral pH and fluoresces brightly in acidic environments, we used it to monitor internalization of α-syn PFF as previously described (*11*). We treated WT or Aplp1 knockout (*Aplp1*^−/−^) primary cortical neurons at 12-14 days *in vitro* with α-syn-pHrodo PFF and assessed the internalization via live-cell imaging. There is endocytosis of the α-syn-pHrodo PFF in WT neurons, whereas the internalization of α-syn-pHrodo PFF was significantly reduced in *Aplp1^−/−^* neurons (Fig. 1G and fig. S2A). Overexpression of Aplp1 by lentivirus transduction significantly increased the endocytosis of α-syn-pHrodo PFF in WT neurons, and re-expression of Aplp1 in *Aplp1^−/−^* neurons restored the endocytosis of α-syn-pHrodo PFF in *Aplp1^−/−^* neurons (Fig. 1G and fig. S2A). Isolation of the endosome fraction via differential centrifugation indicates that α-syn-biotin PFF treatment of cortical neurons led to enrichment of α-syn-biotin PFF in the endosomal fraction in WT neurons, and a significant reduction in *Aplp1^−/−^* neurons (Fig. 1H). Lentiviral transduction of Aplp1 significantly enhanced the amount of α-syn-biotin PFF in the endosomal fraction of WT neurons and *Aplp1^−/−^* neurons (Fig. 1H).

To further support that Aplp1 mediates the endocytosis of α-syn-biotin PFF into endosomes, the intensity of co-localized α-syn-biotin PFF with Rab7 (endosome marker) in WT and *Aplp1^−/−^* cortical neurons was measured. α-Syn-biotin PFF co-localized with Rab7 in WT neurons, whereas in *Aplp1^−/−^* neurons, there were less α-syn-biotin PFF co-localized with Rab7 (Fig. 1I and fig. S2B). Overexpression of Aplp1 in WT or *Aplp1^−/−^* neurons significantly increased the intensity of α-syn-biotin PFF co-localizing with Rab7 (Fig. 1I and fig. S2B). To determine whether Aplp1 also effects the uptake of α-syn-biotin PFF in neurites, the intensity of α-syn-biotin PFF that co-localized with the neuronal marker, microtubule associated protein 2 (MAP2), in neurites was assessed. Similar to observations in the soma, the amount of α-syn-biotin PFF in the MAP2 positive neurites was reduced in *Aplp1^−/−^* neurons compared to WT neurons (fig. S2C).

While overexpression of Aplp1 increased the colocalization of α-syn-biotin PFF with Rab7, overexpression of App or Aplp2 did not increase the intensity of co-localization, compared to control neurons (Fig. 1J). Next, we studied the relationship between the deletion mutants of Aplp1 and intensity of the co-localization of α-syn-biotin PFF with Rab7. While the E1 domain deletion mutant significantly decreases the intensity of co-localization compared to full-length Aplp1 (Fig. 1J), the AcD+E2 domain and ICD deletion mutants do not significantly reduce the intensity of co-localization, compared to full-length Aplp1 (Fig. 1J). These data show that E1 domain of Aplp1 is involved in the endocytosis of α-syn-biotin PFF, but not AcD+E2 or ICD. Furthermore, we investigated the endocytosis of α-syn-biotin PFF by transfecting the Aplp1 chimeras into neurons, and found that both Aplp1(E1)-Aplp2 and Aplp1(E1)-App fail to increase the intensity of the co-localization (Fig. 1K). To exclude the possibility that deletion of Aplp1 causes a generalized defect in uptake, we studied the internalization of latex beads. There was no significant difference in the internalization of latex beads between WT and *Aplp1^−/−^* neurons (fig. S2D). These results show that Aplp1 is involved the endocytosis of α-syn-biotin PFF into neurons and that the E1 domains of Aplp1 is essential for the endocytosis.

### Aplp1 induces α-syn pathology and neurotoxicity

We then asked whether knocking out Aplp1 prevents the pathology induced by α-syn PFF. We administered α-syn PFF or PBS, to WT and *Aplp1^−/−^* cortical neurons at 7 days *in vitro*, and assessed α-syn pathology after ten days. In WT neurons treated with α-syn PFF, the level of phosphorylated serine 129 α-syn (pS129) was significantly increased compared to WT neurons treated with PBS. In *Aplp1^−/−^* neurons treated with α-syn PFF, the level of pS129 was significantly reduced compared to WT neurons treated with α-syn PFF (Fig. 2A and 2B). Overexpression of Aplp1 via lentiviral transduction significantly increased the level of pS129 in both WT neurons and in *Aplp1^−/−^* neurons compared to lentiviral control virus transduction (Fig. 2A and 2B).

**Fig. 2.**
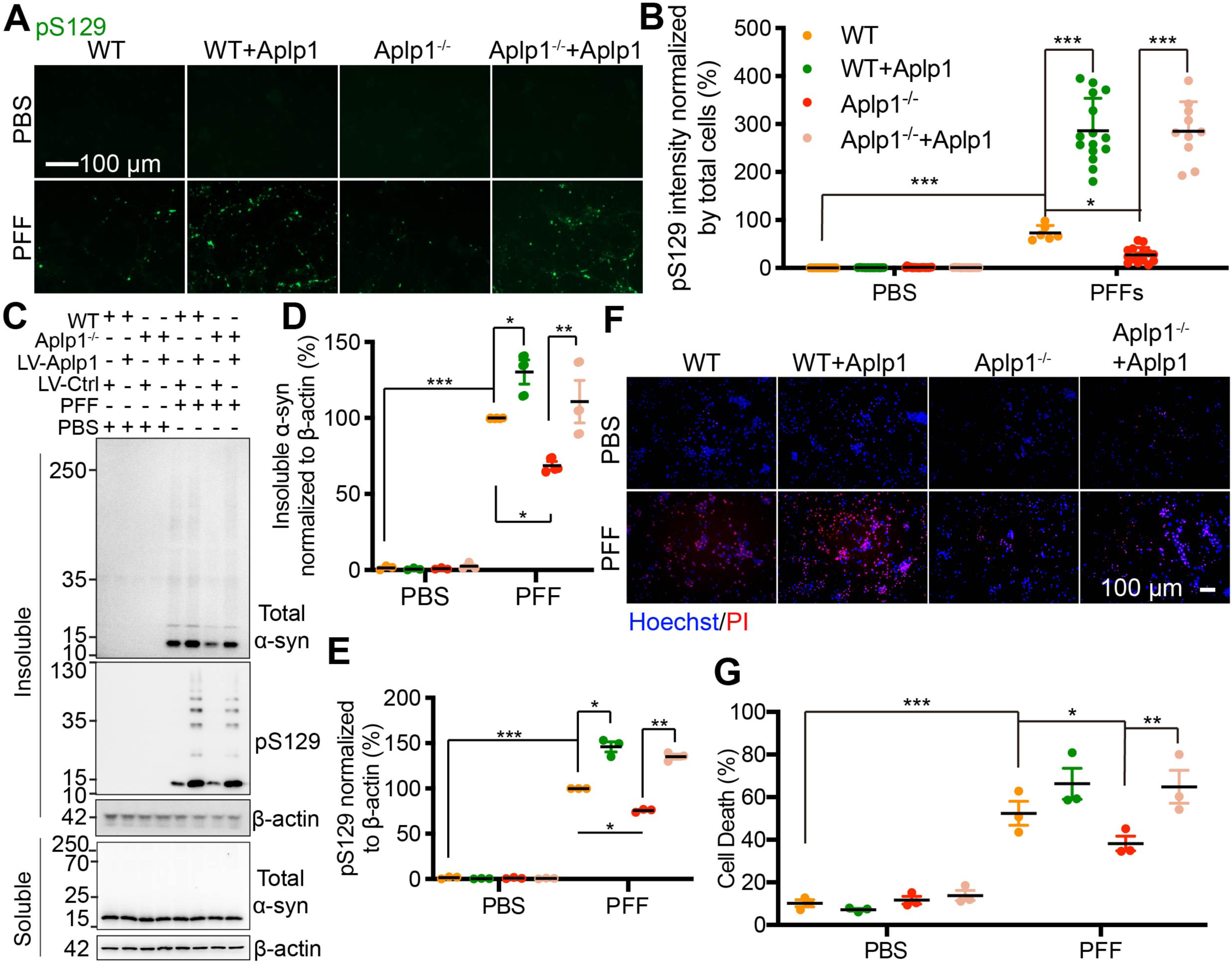
α-Syn PFF-induced pathology and neurotoxicity is reduced by deletion of Aplp1 *in vitro*. **(A)** Immunostaining of anti-pS129 in WT and Aplp1^−/−^ primary cortical neurons by Aplp1 and control lentivirus transduction, with the administration of α-syn PFF and PBS. Scale bar, 100 μm. **(B)** Quantification of (A). *n* = 3 independent experiments. Data are means ± SEM. **(C, D, and E)** Immunoblots and quantification analysis in WT and Aplp1^−/−^ neuron lysates of insoluble α-syn, pS129, soluble α-syn, and β-actin. WT and Aplp1^−/−^ neuron lysates were sequentially extracted in 1% TX-100 (TX-soluble) followed by 2% SDS (TX-insoluble) 14 days after α-syn PFF treatment. α-Syn PFF recruited endogenous α-syn into TX-insoluble and hyper phosphorylated aggregates, which was ameliorated by deletion of Aplp1. *n* = 3 independent experiments. Data are the means ± SEM. **(F)** Aplp1 mediates α-syn PFF toxicity as evaluated by Hoechst and propidium iodide (PI) staining. α-Syn PFF was added into 7 days *in vitro* WT, WT+Aplp1, Aplp1^−/−^, Aplp1^−/−^+Aplp1 primary cortical neurons. The toxicity assay was performed 14-18 days after α-syn PFF treatment. PBS is the negative control. Scale bar, 100 μm. **(G)** Quantification of (F). *n* = 3 independent experiments. Data are as means ± SEM. **(B, D, E, G)** one-way ANOVA followed with Tukey’s correction; * *P* < 0.05, ** *P* < 0.01, *** *P* < 0.001.

Two weeks after α-syn PFF were administered to WT and *Aplp1^−/−^* cortical neurons, we examined α-syn and pS129 levels from lysates sequentially extracted in 1% Triton X-100 (TX-soluble) and 2% SDS (TX-insoluble). In WT neuronal lysates, α-syn PFF led to substantial insoluble α-syn and pS129, whereas in *Aplp1^−/−^* neuronal lysates there was a significant reduction in insoluble α-syn and pS129 (Fig. 2C–2E). Overexpression of Aplp1 via lentiviral transduction in WT and *Aplp1^−/−^* neurons increased the amount of insoluble α-syn and pS129 levels compared to lentiviral control virus transduction (Fig. 2C–2E).

14-18 days after α-syn PFF were administered to WT and *Aplp1^−/−^* cortical neurons, we assessed neurotoxicity with propidium iodide (PI) and Hoechst staining. In WT neurons, α-syn PFF induced substantial neurotoxicity, compared to PBS-treated WT neurons (Fig. 2F and 2G). Depletion of Aplp1 significantly decreased cell death, whereas overexpression of Aplp1 via lentiviral transduction significantly increased the cell death in WT and *Aplp1^−/−^* neurons (Fig. 2F and 2G). Taken together, Aplp1 can drive α-syn PFF-induced pathology and neurotoxicity, and genetic deletion of Aplp1 can significantly reduce α-syn pathology and neurotoxicity.

### Aplp1 and Lag3 bind to each other

Since α-syn PFF binds to both Aplp1 and Lag3(*11*), we hypothesized that Aplp1 and Lag3 bind to each other and work together to facilitate the uptake of α-syn PFF and the subsequent transmission of pathologic α-syn via an Aplp1-Lag3 complex. To test this hypothesis, we investigated the interaction between Aplp1 and Lag3.

Both Aplp1 (*22*) and Lag3 (*11*) proteins are expressed in neurons indicating that they have the potential to interact. Since Lag3 mRNA is enriched in the brain (*24*) and microglia (*25–27*), we further explored the expression of Lag3 in neurons. A Lag3 Loxp reporter line with a YFP (yellow fluorescent protein) signal knocked into the Lag3 locus (*Lag3*^L/L-YFP^) was utilized to determine the cellular localization of Lag3 (*28*). In *Lag3*^L/L-YFP^ mice, YFP marks all cells that express Lag3. Due to nature of gene targeting, YFP is expressed in the cytosol of cells that normally express Lag3 (*28*). The immunoreactivity of anti-GFP, which recognizes YFP co-localized with MAP2 immunostaining in the spinal cord, olfactory bulb, striatum, substantia nigra pars compacta, hippocampus, cortex, cerebellum and brainstem (fig. S3A). The lack of immunoreactivity of anti-GFP in WT mice indicates that the anti-GFP immunoreactivity is specific in *Lag3*^L/L-YFP^ mice (fig. S3B). In addition, anti-GFP colocalizes with IBA1 immunostaining, but not GFAP immunostaining (fig. S3B) consistent with the prior finding that *Lag3* mRNA is in microglia. Immunoreactivity of anti-Aplp1 and anti-Lag3 were observed in the neurons in the cortex of WT mice (fig. S3C). In mice lacking Aplp1 and Lag3 (see fig. S4), immunoreactivity for Aplp1 and Lag3 is absent (fig. S3C). These results confirm that Aplp1 and Lag3 are co-expressed in neurons.

Co-Immunoprecipitation (co-IP) studies showed that Lag3 pulls down Aplp1 by anti-Lag3 410C9 immunoprecipitation in WT mouse brain lysates, but not in *Lag3*^−/−^ lysates (Fig. 3A). Conversely, Aplp1 pulls down Lag3 by anti-Aplp1 CT11 immunoprecipitation in WT mouse brain lysates, but not in *Aplp1*^−/−^ lysates (Fig. 3B). We performed a proximity ligation assay (PLA) (*29, 30*) and the results showed that Aplp1 and Lag3 interact with each other in the cortex of WT mice, but not in *Aplp1*^−/−^*/Lag3*^−/−^ mice (Fig. 3C). To identify the Lag3-binding domain in Aplp1, we conducted co-immunoprecipitation experiments with the FLAG-Aplp1 deletion mutants described previously (see fig. S1C-E). The results showed that the E1 domain of FLAG-Aplp1 is involved in the interaction with Lag3-Myc (Fig. 3D). As Lag3-Myc binding to FLAG-Aplp1 is eliminated by deletion of the E1 domain of FLAG-Aplp1, we studied two additional deletion mutants of the E1 subdomain (ΔGFLD and ΔCuBD). The ΔGFLD mutant substantially reduced Lag3-Myc binding to FLAG-Aplp1, while the ΔCuBD mutant reduced binding to a lesser extent (Fig. 3E), suggesting that GFLD subdomain in the E1 domain of Aplp1 (residues 48-154) is the major subdomain responsible for the Lag3 interaction. Similarly, the Aplp1-binding domain in Lag3 was determined via co-IP experiments. The D2 and D3 domains of Lag3-Myc are essential for the FLAG-Aplp1 interaction (Fig. 3F). Confirmation that the D2 and D3 deletion constructs of Lag3-Myc reach the cell membrane is provided in fig. S1.

**Fig. 3.**
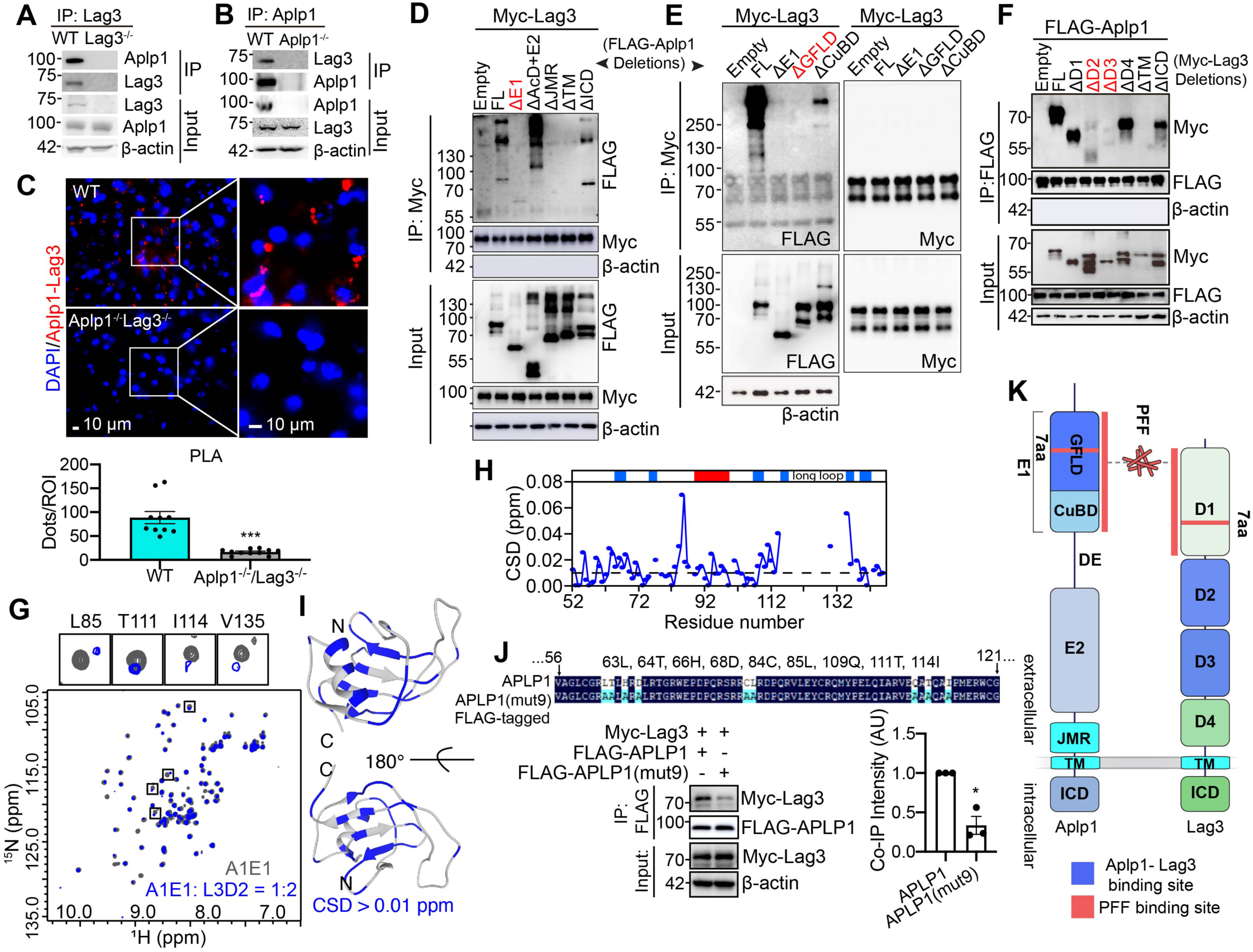
Aplp1 and Lag3 bind to each other. **(A)** Lag3 pulls down Aplp1 by anti-Lag3 410C9 immunoprecipitation in WT mouse brain lysates, but not in *Lag3^−/−^* lysates. **(B)** Aplp1 pulls down Lag3 by anti-Aplp1 CT11 immunoprecipitation in WT mouse brain lysates, but not in *Aplp1^−/−^* lysates. **(C)** Proximity ligation assay (PLA) staining images and quantification for the Aplp1 and Lag3 interaction in the cortex region of WT and *Aplp1^−/−^/Lag3^−/−^* mouse brain sections. The red dots around the nucleus (blue) represent the PLA signal for the Aplp1 and Lag3 interaction. Scale bars, 10 μm. The analysis was obtained from WT mice (2 mice, 11 images) and *Aplp1^−/−^/Lag3^−/−^* mice (2 mice, 11 images). Student’s *t*-test, \*\**P* < 0.01. **(D and E)** Mapping of the Lag3-binding domains in Aplp1. HEK293FT cells were transfected with full-length (FL) or deletion mutants of FLAG-Aplp1, and Myc-Lag3 for co-IP experiments. The GFLD subdomain in the E1 domain of Aplp1 is the major subdomain responsible for the Lag3 interaction. **(F)** Mapping of the Aplp1-binding domains in Lag3. HEK293FT cells were transfected with full-length (FL), deletion mutants of Myc-Lag3, and FLAG-Aplp1 for co-IP experiments. **(G, H, and J)** Identification of the interface of A1E1 (E1 domain of APLP1) binding to L3D2 (D2 domain of LAG3). **(G)** Overlay of the 2D ^1^H-^15^N HSQC spectra of A1E1 alone (grey) and in the presence of 2 molar folds of L3D2 (blue). Four residues with significant CSDs (> 0.03 ppm) are highlighted and enlarged in the black boxes. **(H)** Histogram of the calculated chemical shift deviations (CSDs) of A1E1 in the presence of L3D2 at a molar ratio of 1:2 (A1E1/L3D2). The domain organization of A1E1 is indicated on the top, with blue boxes indicating the *β*-strands and the red box indicating the *α*-helix. A dashed line was drawn to highlight the residues with CSDs > 0.01 ppm. **(I)** The 37 residues with large CSDs (> 0.01 ppm) upon L3D2 titration are highlighted in blue on the ribbon diagram of A1E1 modelled structure. **(J)** Validation of the NMR results. We substituted nine residues of APLP1 with alanine to generate FLAG-APLP1(mut9), and performed the co-IP experiment to assess the APLP1-Lag3 interaction. *n* = 3 independent experiments. Data are the means ± SEM, Student’s *t*-test; \**P* < 0.05. **(K)** The scheme for the interaction among Aplp1, Lag3 and α-syn PFF.

We next sought to investigate the structural basis for the interaction between the extracellular domains of APLP1 and Lag3, by using nuclear magnetic resonance (NMR) spectroscopy. Since our cellular data suggests that the E1 domain of Aplp1 binds to Lag3 via its D2 and D3 domains (Fig. 3D–3F), we purified recombinant E1 domain of APLP1 (A1E1), and D2 and D3 domains of Lag3 (L3D2 and L3D3), and performed NMR titration experiments to study their interaction. The titration of L3D2 to A1E1 results in significant, but relatively small chemical shift deviations (CSDs, > 0.01 ppm) of 37 residues of A1E1 (Fig. 3G and 3H), implying direct and weak binding of the two components. When the 37 residues were mapped to the structural model of A1E1 (Fig. 3I), we found that they are mainly located across the central *β*-sheet of A1E1. These results demonstrate that L3D2 directly interacts with A1E1 via binding to its *β*-sheet region. Of note, five residues (R90, V91, Y94, Q97, M98) located in the *α*-helix region of A1E1 (residue 90–98) also showed CSDs > 0.01 ppm. However, these residues face to the inner side of the structure, implying that their CSDs may result from the conformational changes of its neighboring *β*-sheet, rather than the direct binding of L3D2 to the *α*-helix region of A1E1.

We further examined the interaction between L3D3 and A1E1 by NMR titration. In comparison to L3D2, titration of L3D3 to A1E1 resulted in CSDs (> 0.01 ppm) of 5 residues of A1E1 (C84, L85, T111, I114, V135), suggesting that L3D3 is capable of binding to A1E1, but the binding affinity is lower than that of L3D2 to A1E1 (fig. S3D). Based on the NMR results, we substituted nine residues of APLP1 with alanine to generate FLAG-APLP1(mut9), and performed the co-IP experiment to assess the APLP1-Lag3 interaction (Fig. 3J). Co-IP results showed that the substitution significantly decreases the interaction between APLP1 and Lag3 (Fig. 3J). Taken together, these results demonstrate that A1E1 directly interacts with both D2 and D3 of Lag3, to form the Aplp1-Lag3 complex (Fig. 3K).

### The Aplp1 and Lag3 complex role in binding and endocytosis of α-syn PFF

Double knockout of Aplp1 and Lag3 (*Aplp1*^−/−^*/Lag3*^−/−^) mice were generated to determine the potential consequences of the interaction between Aplp1 and Lag3 on α-syn PFF-induced neurodegeneration (fig. S4A and S4B). α-Syn-biotin PFF-binding was reduced in both *Aplp1*^−/−^ and *Lag3*^−/−^ mice cortical neuron cultures (fig. S4C). α-Syn-biotin PFF-binding was reduced even further in *Aplp1*^−/−^*/Lag3*^−/−^ cortical cultures (fig. S4C). Together Aplp1 and Lag3 account for greater than 40% of the binding of α-syn-biotin PFF to cortical neurons (Fig. 4A). *Aplp1*^−/−^*/Lag3*^−/−^ cortical neurons were transduced with Aplp1, Lag3, or Aplp1 + Lag3 by lentivirus and α-syn-biotin PFF binding was assessed. Individual overexpression of Aplp1 or Lag3 leads to increased binding over baseline of α-syn-biotin PFF to *Aplp1*^−/−^*/Lag3*^−/−^ cortical neurons (Fig. 4B and fig. S4C). Re-expression of Aplp1 and Lag3 together in *Aplp1*^−/−^*/Lag3*^−/−^ cortical neurons led to a 70% enhancement of binding that is greater than the sum of α-syn-biotin PFF binding to Aplp1 and Lag3 when individually expressed in *Aplp1*^−/−^*/Lag3*^−/−^ cortical neurons (Fig. 4B and fig. S4D).

**Fig. 4.**
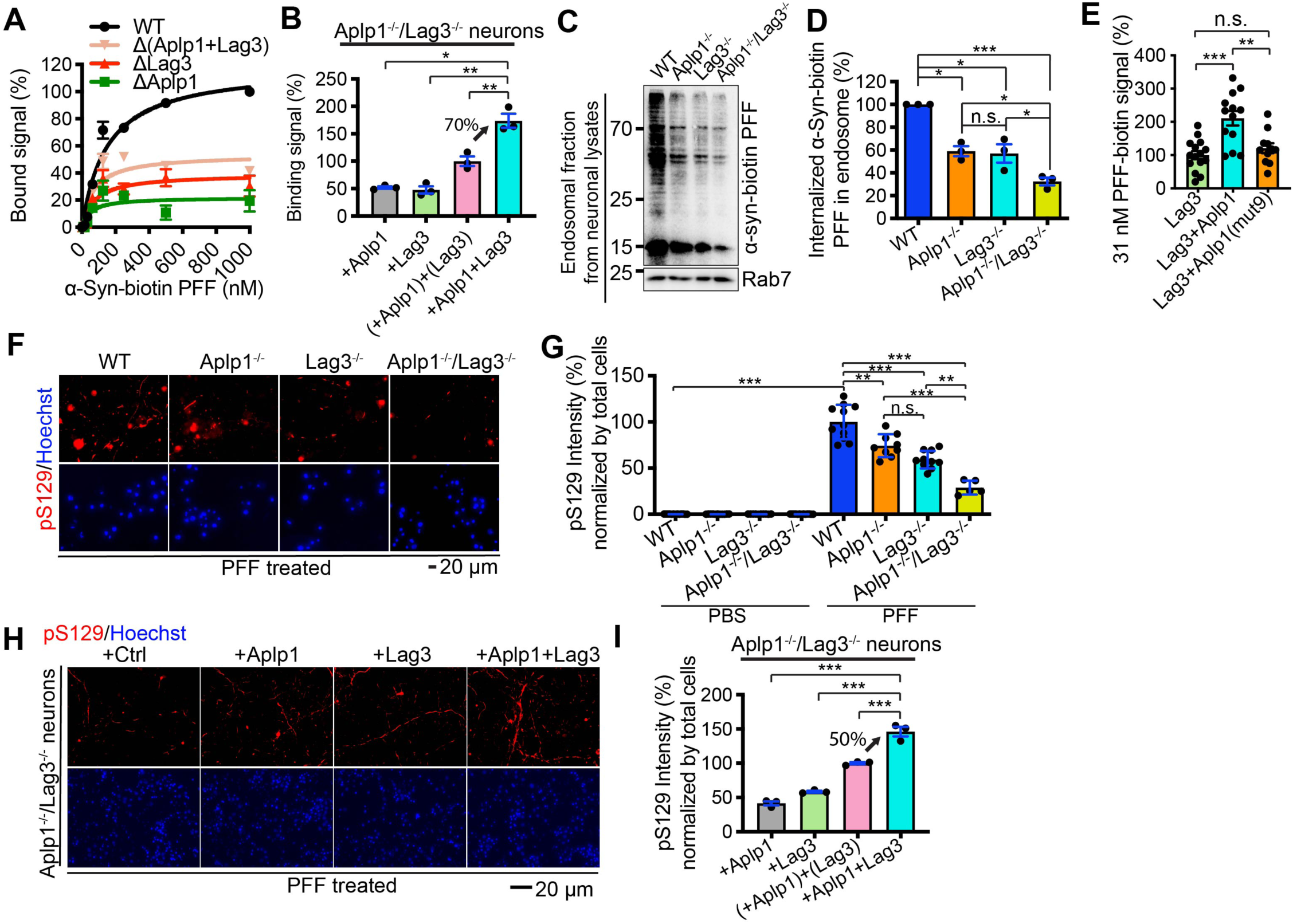
The role of Aplp1-Lag3 complex in mediating the binding, endocytosis of α-syn PFF, and subsequent pathology. **(A)** Aplp1 and Lag3 account for greater than 40% of the binding of α-syn-biotin PFF to cortical neurons analyzed by Fig. S4C. **(B)** Re-expression of Aplp1 and Lag3 together in *Aplp1*^−/−^*/Lag3*^−/−^ cortical neurons led to 70% enhanced binding that is greater than the sum of α-syn-biotin PFF binding to Aplp1 and Lag3 alone. Refer to Fig. S4D. One-way ANOVA followed by Dunnett’s correction. **(C)** Immunoblots and **(B)** quantification analysis of endosomal fractions isolated from WT, *Aplp1^−/−^*, *Lag3^−/−^*, and *Aplp1^−/−^/Lag3^−/−^* cortical neurons. Knocking out the Aplp1-Lag3 complex (Aplp1^−/−^/Lag3^−/−^) induced a significant decrease (∼70%) in the amount of internalized α-syn-biotin PFF compared to WT neurons, and it is also significantly less than the depletion of Aplp1 or Lag3 alone. Rab7 is the loading control. One-way ANOVA followed by Tukey’s correction. **(E)** Aplp1 co-expression by transfection increased the uptake of α-syn-biotin in Lag3-transfected SH-SY5Y cells, and Aplp1(mut9) expression failed to increase the uptake. Cells were incubated with 31 nM α-syn-biotin for 2 hr at 37° C. The biotin signal was normalized using the intensity of Lag3. Lag3 (15 cells), Lag3+Aplp1 (14 cells), Lag3+Aplp1(mut9) (11 cells). Data are the means ± SEM. \*\**P* < 0.01, \*\*\**P* < 0.001; n.s., not significant; *n* = 3 independent experiments. **(F and G)** Deletion of the Aplp1-Lag3 complex significantly decreased (70%) pS129 immunostaining induced by α-syn PFF, compared to Aplp1^−/−^, Lag3^−/−^, or WT neurons. Scale bar, 20 μm. Data are the means ± SEM. **(H and I)** Immunostaining of anti-pS129 in *Aplp1*^−/−^/*Lag3*^−/−^ neurons, treated with α-syn PFF, transduced with Aplp1, Lag3, or Aplp1+Lag3. Scale bar, 20 μm. Data are the means ± SEM. *n* = 3 individual experiments, one-way ANOVA with Turkey’s correction. \*\**P* < 0.01, \*\*\**P* < 0.001, n.s., no significance.

We examined the internalization of α-syn-biotin PFF in the endosome-enriched fractions isolated from WT, *Aplp1*^−/−^, *Lag3*^−/−^, and *Aplp1*^−/−^*/Lag3*^−/−^ cortical neurons. There was significantly less internalized α-syn-biotin PFF in *Aplp1*^−/−^ or *Lag3*^−/−^ neurons than in WT neurons (Fig. 4C and 4D). Knocking out both Aplp1 and Lag3 induced a significant decrease (∼70%) in the amount of internalized α-syn-biotin PFF compared to WT neurons, which was also significantly less than the depletion of Aplp1 or Lag3 alone (Fig. 4C and 4D). To exclude the possibility that double deletion of Aplp1 and Lag3 causes a defect in endocytosis, we studied the internalization of latex beads. There was no significant difference in the internalization of latex beads between WT, *Aplp1*^−/−^, *Lag3*^−/−^, and *Aplp1*^−/−^*/Lag3*^−/−^ cortical neurons (see fig. S2D). We administered α-syn-biotin PFF into Lag3-transfected cell cultures, and found that Aplp1 co-expression increased the uptake signal of α-syn-biotin PFF (Fig. 4E). Aplp1(mut9) expression failed to increase the uptake of α-syn-biotin PFF in Lag3-transfected cells (Fig. 4E). Of note there is no appreciable difference in the signal of α-syn-biotin PFF uptake in Aplp1 versus Aplp1(mut9) transfected cells (Fig.S4E) suggesting the Aplp1(mut9) itself does not appreciably interfere with α-syn-biotin PFF uptake. Taken together these results indicate the depletion of both Aplp1 and Lag3 significantly reduce the binding and the internalization of α-syn-biotin PFF, and Aplp1 and Lag3 form a complex that significantly enhances the binding and uptake of α-syn-biotin PFF.

### The role of the Aplp1-Lag3 complex in α-syn pathology, transmission and neurotoxicity

To assess the role of the Aplp1-Lag3 complex in α-syn pathology following α-syn PFF administration, α-syn PFF were incubated for 12 days in WT, *Aplp1*^−/−^, *Lag3*^−/−^, and *Aplp1*^−/−^*/Lag3*^−/−^ primary cortical neurons. Depletion of both Aplp1 and Lag3 significantly decreased pS129 immunoreactivity by 70% that was induced by α-syn PFF compared to WT neurons. This reduction is significantly less than individual Aplp1^−/−^ or Lag3^−/−^ neurons (Fig. 4F and 4G). The levels of pS129 were assessed in *Aplp1*^−/−^*/Lag3*^−/−^ neurons transduced with Aplp1, Lag3 or Aplp1 + Lag3 via lentivirus following α-syn PFF treatment. Re-expression of Aplp1 and Lag3 together in *Aplp1*^−/−^*/Lag3*^−/−^ cortical neurons led to a greater than 50% enhanced pS129 intensity that is larger than the sum of the individual contribution of the increased pS129 intensity when Aplp1 and Lag3 re-expressed individually in *Aplp1*^−/−^*/Lag3*^−/−^ neurons (Fig. 4H and 4I).

A microfluidic neuronal culture device with three chambers, which are connected in tandem by two series of microgrooves (TCND1000, Xona Microfluidics) was used to examine transmission of pathologic α-syn in WT, *Aplp1*^−/−^, *Lag3*^−/−^, and *Aplp1*^−/−^*/Lag3*^−/−^ cortical cultures (Fig. 5A). Transmission of pathologic α-syn was monitored by pS129 immunoreactivity as previously described^11^ (Fig. 5B). Four groups of neuron combinations were set up in successive chambers (1)-(2)-(3): (WT)-(WT)-(WT), (WT)-(*Aplp1*^−/−^)-(WT), (WT)-(*Lag3*^−/−^)-(WT), and (WT)-(*Aplp1*^−/−^*/Lag3*^−/−^)-(WT). There was an equivalent pS129 level in chamber 1 (C1) in *Aplp1*^−/−^, *Lag3*^−/−^, and *Aplp1*^−/−^*/Lag3*^−/−^ cultures, which indicates that the pathologic α-syn levels in the WT neurons are at the same level at the initiation of the experiment (Fig. 5C). *Aplp1*^−/−^, *Lag3*^−/−^, and *Aplp1*^−/−^*/Lag3*^−/−^ significantly blocked the transmission of pathologic α-syn to chamber 2 (C2) and chamber 3 (C3) (Fig. 5D and 5E), suggesting that both Aplp1 and Lag3 are required for the transmission of pathologic α-syn.

**Fig. 5.**
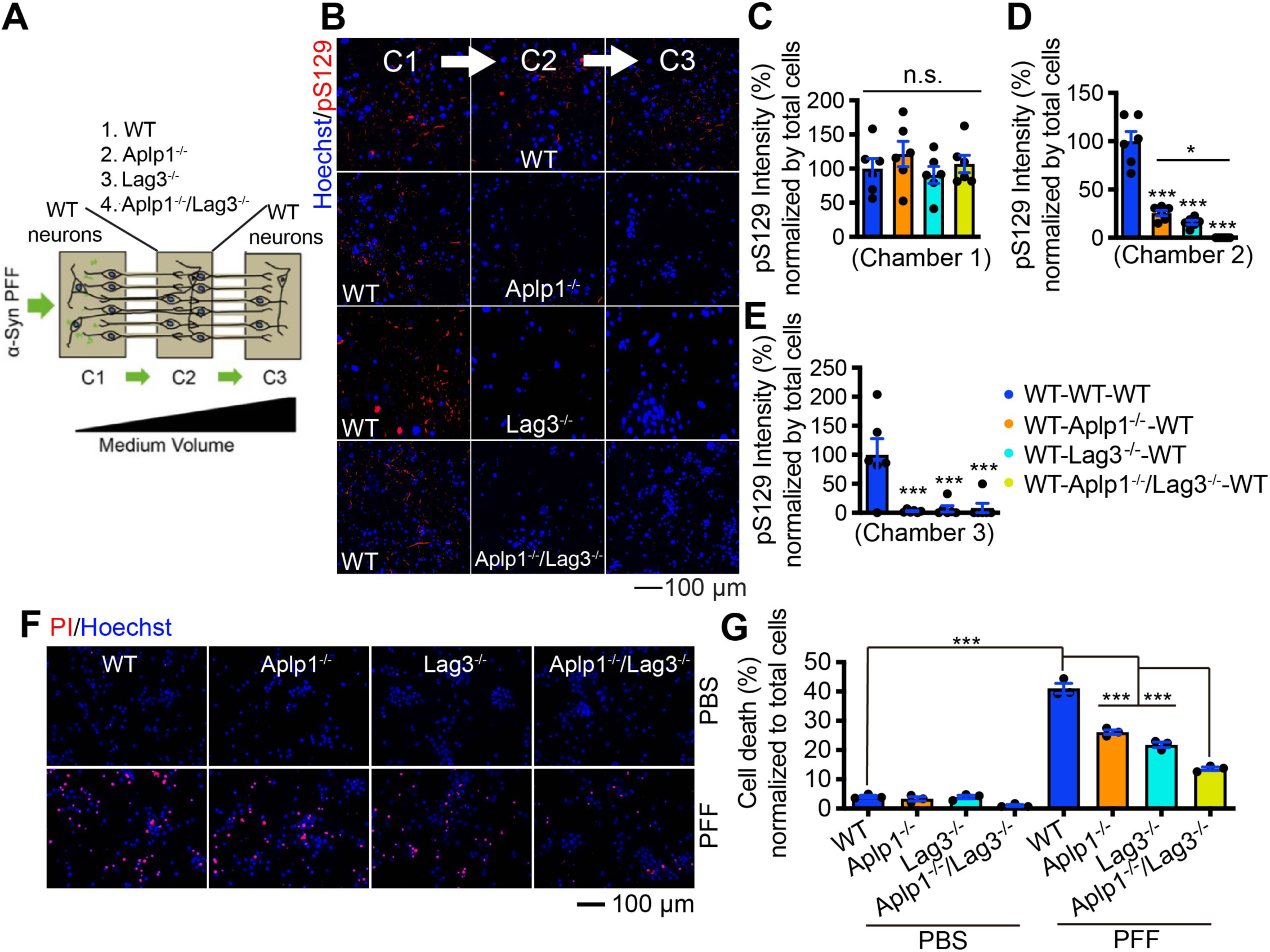
The role of Aplp1-Lag3 complex in pathologic α-syn transmission and neurotoxicity *in vitro*. **(A)** Schematic of microfluidic neuron device with three chambers. α-Syn transmission from chamber 1 (C1) to chamber 2 (C2), then to chamber 3 (C3) 14 days after addition of α-syn PFF in C1. The different combinations of neurons tested in C2, listed as C1-(C2)-C3, are WT-(WT)-WT, WT-(Aplp1^−/−^)-WT, WT-(Lag3^−/−^)-WT, WT-(Aplp1^−/−^/Lag3^−/−^)-WT. **(B**–**E)** Immunostaining images of pS129 in the transmission. Scale bar, 100 μm. Values are means ± SEM. **(F and G)** PI/Hoechst staining for cell death in WT, *Aplp1*^−/−^, *Lag3*^−/−^, and *Aplp1*^−/−^/*Lag3*^−/−^ cortical neuron cultures, treated with α-syn PFF. Scale bar, 100 μm. Deletion of the Aplp1-Lag3 complex exhibited significantly less cell death than in *Aplp1*^−/−^, *Lag3*^−/−^ or WT cultures, treated with α-syn PFF. Data are the means ± SEM. *n* = 3 individual experiments, one-way ANOVA with Turkey’s correction. \**P* < 0.05, \*\**P* < 0.01, \*\*\**P* < 0.001, n.s., no significance.

PI/Hoechst staining was used to assess cell death in cortical neuron cultures of WT, *Aplp1*^−/−^, *Lag3*^−/−^, and *Aplp1*^−/−^*/Lag3*^−/−^ following administration of α-syn PFF. *Aplp1*^−/−^*/Lag3*^−/−^ cultures exhibited significantly less cell death than *Aplp1^−/−^*, *Lag3^−/−^* or WT cultures (Fig. 5F and 5G).

### Roles of Aplp1 and the Aplp1-Lag3 complex in mediating α-syn PFF-induced neurodegeneration *in vivo*

To determine the roles of Aplp1 and the Aplp1-Lag3 complex contributions to neurodegeneration induced by α-syn PFF *in vivo*, we stereotactically injected the same amount of α-syn PFF (5 μg) into the dorsal striatum of *Aplp1*^−/−^ and *Aplp1*^−/−^*/Lag3*^−/−^. As previously described in WT mice at 180 days after injection (*10, 11*), there was a significant loss of dopamine (DA) neurons as detected by stereologic counting of tyrosine hydroxylase (TH)- and Nissl-positive neurons in the substantia nigra pars compacta (SNpc) (Fig. 6A and 6B). There was a moderate rescue of DA neurons in α-syn PFF-injected *Aplp1*^−/−^ mice (Fig. 6A and 6B). Depletion of App (*App*^−/−^) had no effect on DA neuron loss (fig. S5A and S5B) providing specificity to the role of Aplp1 in contributing to pathologic α-syn-induced neurodegeneration. In α-syn PFF-injected *Aplp1*^−/−^*/Lag3*^−/−^ mice, there was an extensive preservation of DA neurons (Fig. 6A and 6B). In addition, although intrastriatal injection of α-syn PFF induced a decrease in striatal TH-positive immunoreactivity in WT mice, in *Aplp1*^−/−^ mice there was preservation of the striatal TH-positive immunoreactivity and in the *Aplp1*^−/−^*/Lag3*^−/−^ mice there was significantly greater preservation of TH-positive immunoreactivity (fig. S5C and S5D). High-performance liquid chromatography-electrochemical detection (HPLC-ECD) analysis demonstrated a significant reduction in the levels of DA and its metabolites homovanillic acid (HVA), 3,4-dihydroxyphenylacetic acid (DOPAC), and 3MT in WT mice by α-syn PFF (Fig. 6C and fig. S5E–G). Depletion of Aplp1 mildly alleviated the DA deficits but the results are not significantly different (Fig. 6C and fig. S5E–G). The DA deficits were eliminated in the *Aplp1*^−/−^*/Lag3*^−/−^ mice (Fig. 6C and fig. S5E–G). Immunoreactivity for pS129 was monitored in the SNpc TH-positive neurons 180 days after α-syn PFF injection. As previously described (*10, 11*), we observed substantial pS129 immunostaining in WT SNpc TH-positive neurons (Fig. 6D and 6E). In contrast, in *Aplp1*^−/−^ SNpc TH-positive neurons, pS129 immunostaining was significantly reduced by approximately 60%, and depletion of both Aplp1 and Lag3 (*Aplp1*^−/−^*/Lag3*^−/−^ mice) significantly reduced the pS129 immunostaining by approximately 90% (Fig. 6D and 6E).

**Fig. 6.**
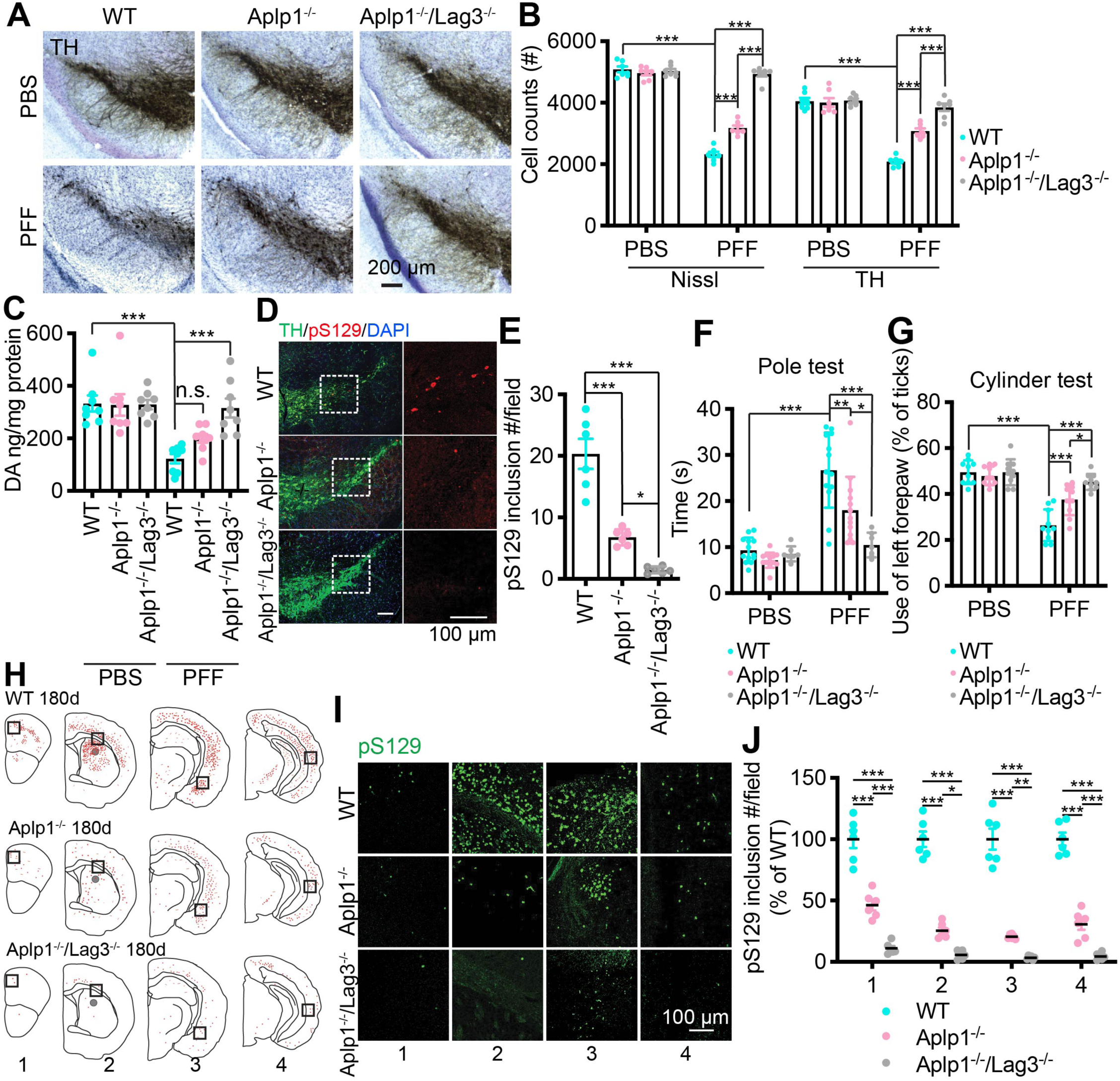
Roles of Aplp1 and the Aplp1-Lag3 complex in mediating α-syn PFF-induced neurodegeneration *in vivo*. **(A)** Representative TH (tyrosine hydroxylase) immunohistochemistry and Nissl staining images of dopamine (DA) neurons in the SNpc of α-syn PFF-injected hemisphere in the WT, *Aplp1^−/−^*, and *Aplp1^−/−^/Lag3^−/−^* mice. **(B)** Stereological counting of the number of TH- and Nissl-positive neurons in the substantia nigra via unbiased stereological analysis after 6 months of α-syn PFF injection in the WT, *Aplp1^−/−^*, and *Aplp1^−/−^/Lag3^−/−^* mice (WT: *n* = 7; *Aplp1^−/−^*: *n* = 7; *Aplp1^−/−^/Lag3^−/−^*: *n* = 7). Data are the means ± SEM, one-way ANOVA with Tukey’s correction; \*\*\**P* < 0.001. **(C)** DA concentrations in the striatum of α-syn PFF-injected mice and PBS-injected controls measured at 180 days by means of HPLC. Data are the means ± SEM, *n* = 8 mice per each group, one-way ANOVA with Dunnett’s correction. **(D and E)** Representative pS129 positive inclusions in the substantia nigra of WT, *Aplp1^−/−^*, and *Aplp1^−/−^/Lag3^−/−^* mice. The pS29 immunopositive neurons were quantified in the substantia nigra of WT, *Aplp1^−/−^*, and *Aplp1^−/−^/Lag3^−/−^* mice (*n* = 6, each group). **(F and G)** Assessments of the behavioral deficits measured by the pole test (WT: *n* = 13; *Aplp1^−/−^*: *n* = 13; *Aplp1^−/−^/Lag3^−/−^*: *n* = 10, each group) and cylinder test (*n* = 10, each group). **(H)** Distribution of LB/LN-like pathology in the CNS of α-syn PFF-injected hemisphere of WT, *Aplp1^−/−^*, and *Aplp1^−/−^/Lag3^−/−^* mice (pS129 positive neuron, red dots; pS129 positive neurites, red lines). **(I and J)** Representative images of LB/LN-like pathology (the black box in panel H) and the quantification of pS129 intensity (green) from each coronal section (1–4) stained with pS129 by ImageJ. The *in vivo* work was performed in a double-blinded fashion. Data are the means ± SEM. Statistical significance was determined by using one-way ANOVA followed with Tukey’s correction, \**P* < 0.05, \*\**P* < 0.01, \*\*\**P* < 0.001, n.s., not significant.

α-Syn transmission *in vivo* was monitored by assessing the distribution of pS129 α-syn pathology in the coronal brain sections of WT, *Aplp1*^−/−^ and *Aplp1*^−/−^*/Lag3*^−/−^ mice (Fig. 6H–6J). We observed substantial immunoreactivity of pS129 in coronal brain sections of WT mice at 180 days after injection (Fig. 6H–6J). In contrast, in *Aplp1*^−/−^ mice, the immunoreactivity of pS129 was significantly reduced by approximately 60% and depletion of the Aplp1-Lag3 complex significantly blocked the immunoreactivity of pS129 by greater than 90% (Fig. 6H–6J).

WT mice injected with α-syn PFF were significantly impaired in their performance on the pole test and the cylinder test (Fig. 6F and 6G). These behavioral deficits were significantly reduced in the *Aplp1*^−/−^ mice and were eliminated in the *Aplp1*^−/−^*/Lag3*^−/−^ mice (Fig. 6F and 6G).

### Role of Lag3 antibody in blocking α-syn PFF-induced neurodegeneration *in vivo*

Previously we showed that the Lag3 antibody, 410C9, significantly decreased the internalization of α-syn-biotin PFF in WT cortical neurons (*11*). Accordingly, the domain of Lag3 that 410C9 recognizes was assessed to determine whether it could serve as a reagent to examine the role of the interaction of Lag3 with Aplp1 in the internalization of pathologic α-syn. We found that 410C9 recognizes the D3 domain of Lag3 (fig. S6A) (*31*). Since Aplp1 interacts with D3 domain of Lag3 (see Fig. 3F, fig. S3D, and Fig. 3K), we assessed whether 410C9 could disrupt the Aplp1-Lag3 complex. 410C9 significantly disrupts the co-IP of FLAG-Aplp1 and Lag3-Myc in HEK293FT cells (fig. S6B and S6C). α-Syn PFF treatment of the cellular extract of FLAG-Aplp1 and Lag3-Myc transfected HEK293FT cells significantly increased the co-IP of FLAG-Aplp1 and Lag3-Myc (fig. S6D and S6E). We compared the inhibitory effects of 410C9 on the internalization of α-syn-biotin PFF in WT versus *Lag3*^−/−^ cortical neurons. In the endosomal fraction, 410C9 blocked the internalization of α-syn-biotin PFF significantly more in WT cultures than in *Lag3*^−/−^ cultures (fig. S6F and S6G). Both the levels of the endosomal Aplp1 and Lag3 were decreased by 410C9 in WT cortical cultures (fig. S6H). Taken together these results further support the Aplp1-Lag3 complex, and suggest that 410C9 disrupts the Aplp1 and Lag3 complex leading to enhanced blockade of the internalization of α-syn PFF, which is greater than that observed in *Lag3*^−/−^ cultures.

Next, we asked whether 410C9 could block the neurodegeneration induced by intrastriatal injection of α-syn PFF. WT mice were inoculated with α-syn PFF and then were treated with 410C9 or control mouse IgG (mIgG) (10 mg/kg, intraperitoneally [i.p.]), followed by weekly treatment with 410C9 or mIgG for 180 days (a total of 26 treatments) (fig. S7A). To determine whether i.p. administration of 410C9 led to an appreciable antibody concentration within the brain, we measured 410C9 levels in the cerebral spinal fluid (CSF) at 3, 7, and 10 days following a single i.p. injection (10 mg/kg) (*32*). Within 3 days following injection, the CSF levels of 410C9 reached 0.5% of the plasma levels similar to what has been reported with other antibodies that cross the blood brain barrier (*33, 34*), and then persisted for up to 10 days (fig. S7B and S7C). In contrast, the plasma levels of 410C9 significantly decreased at day 7, was not detectable at day 10 (fig. S7D).

To test the efficacy of 410C9 in inhibiting the internalization of α-syn PFF in neurons *in vivo*, WT mice were pre-treated with 410C9 or mIgG (10 mg/kg) for 3 days, followed by intrastriatal injection of α-syn-biotin PFF. We examined the co-localized intensity of α-syn-biotin PFF with Rab7 in neurons with MAP2-positive staining, and found that 410C9 significantly inhibited the co-localization of α-syn-biotin PFF with Rab7 *in vivo* (fig. S7E and S7F).

To test whether 410C9 retards disease progression (DA neuron loss, α-syn pathology, DA levels, and behavioral deficits), we initiated treatment with either 410C9 or mIgG in separate cohorts one day post intrastriatal injection of α-syn PFF or PBS. At 180 days after α-syn PFF injection, stereologic counting of TH- and Nissl-positive neurons in the SNpc revealed a significant loss of DA neurons in WT mice treated with mIgG (Fig. 7A and 7B). In contrast, 410C9 significantly reduced the loss of DA neurons induced by α-syn PFF (Fig. 7A and 7B).

**Fig. 7.**
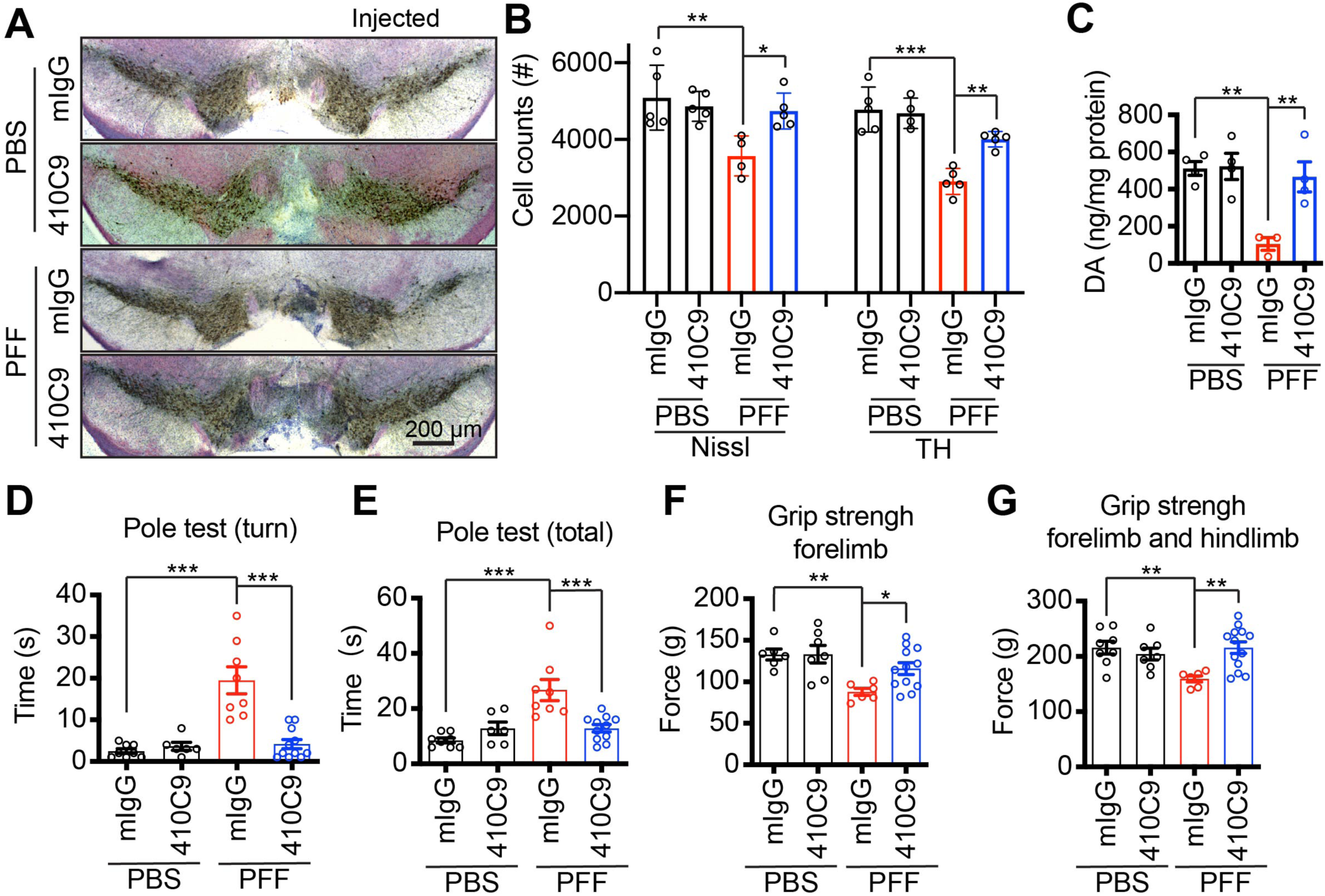
The role of anti-Lag3 in blocking α-syn PFF-induced neurodegeneration *in vivo*. **(A)** Representative images of TH and Nissl staining of SNpc DA neurons of WT mice treated with mIgG or 410C9 at 6-months after intrastriatal injection of α-syn PFF injection. **(B)** Stereology counts of data are the means ± SEM, (mIgG-PBS: *n* = 5; 410C9-PBS: *n* = 5; mIgG-PFF: *n* = 5; 410C9-PFF: *n* = 6). One-way ANOVA with Turkey’s correction. \**P* < 0.05, \*\**P* < 0.01, \*\*\**P* < 0.001. **(C)** DA concentrations in the striatum of α-syn PFF or PBS injected mice treated with 410C9 or mIgG measured at 6-months by means of HPLC-ECD (high performance liquid chromatography-electrochemical detection). **(D–G)** Behavioral deficits were ameliorated by 410C9. Six-months after α-syn PFF injection, the grip strength test and pole test were performed. Behavioral abnormalities in the grip strength test and pole test induced by α-syn PFF were ameliorated by 410C9 treatment (one day after stereotaxic injection). The *in vivo* work was performed in double-blind fashion. Data are the means ± SEM, (mIgG-PBS: *n* = 8; 410C9-PBS: *n* = 6; mIgG-PFF: *n* = 8; 410C9-PFF: *n* = 11). Statistical significance was determined by using one-way ANOVA with Dunnett’s correction; \**P* < 0.05, \*\**P* < 0.01, \*\*\**P* < 0.001.

In PFF-inoculated, mIgG-treated mice, pS129 α-syn was measured in three brain regions: the cortex, amygdala, and SNpc, primarily ipsilateral to the injection site (fig. S7G–S7J). Importantly, 410C9 treatment significantly decreased the level of pS129 α-syn in these brain regions (fig. S7G–S7J). Thus, the simultaneous administration of 410C9 led to reduction in α-syn PFF-induced pS129 α-syn pathology *in vivo*.

Immunoblot analysis demonstrated a significant decrease in TH and the DA transporter (DAT) in the PFF-injected WT mice treated with mIgG (fig. S7K–S7M), compared to the PBS-injected mice treated with mIgG. Importantly, 410C9 treatment significantly increased the expression of TH and DAT (fig. S7K–S7M).

The decrease of DA and its metabolites DOPAC, HVA, and 3MT in WT mice by α-syn PFF, can be prevented by 410C9, as detected by HPLC-ECD (Fig. 7C and fig. S7N–S7P). Furthermore, the behavioral deficits by α-syn PFF in the pole test and grip strength test were alleviated by 410C9 (Fig. 7D–7G).

Taken together these data indicate that Lag3 immunotherapy decreased the spread of pathologic α-syn *in vivo*, ameliorated motor dysfunction mediated by LB pathology, and rescued TH cell loss.

## DISCUSSION

The major findings of the paper are that Aplp1 is a receptor for pathologic α-syn that facilitates pathologic α-syn transmission, and that together with Lag3 it forms a Aplp1-Lag3 complex that contributes to pathologic α-syn transmission and pathogenesis. Double knockout of Aplp1 and Lag3 and or an anti-Lag3 antibody that disrupts the Aplp1 and Lag3 complex almost completely blocks α-syn PFF-induced neurodegeneration and behavioral deficits.

Emerging evidence suggests that misfolded α-syn binds to at least eight transmembrane proteins including heparan sulfate proteoglycans (HSPGs) (*14*), TLR2 (*13, 15*), neurexins (*11, 16, 35*), Na+/K+-ATPase subunit α3 (*16*), Lag3 (*11*), Aplp1 (*11*), FcγRIIb (*12*) and the cellular prion protein (PrPc) (*17*). Of these α-syn binding proteins, HSPGs, Lag3, TLR2, FcγRIIb and PrPc have been reported to be either directly or indirectly involved in the internalization of pathologic α-syn. Our findings indicate that Aplp1 plays a role in the internalization of pathologic α-syn in neurons and contributes to the spread and neurodegeneration by induced by pathologic α-syn. Aplp1 and Lag3 form a complex that accounts for greater than 40% of the neuronal binding and 70% of the neuronal uptake. In addition, the Aplp1-Lag3 complex accounts for the majority (> 90%) of the pathologic spreading and toxicity induced by pathologic α-syn. Our findings that the Lag3 antibody, 410C9 disrupts the interaction of Aplp1 and Lag3 and dramatically protects against the degenerative process set in motion by pathologic α-syn that is equivalent to deleting both Aplp1 and Lag3 supports that notion that the Lag3-Aplp1 complex is important in degenerative process induced by cell-to-cell spread of pathologic α-syn. Thus, other α-syn binding proteins likely either play accessory roles in the endocytosis and toxicity of pathologic α-syn or are involved in other roles. For instance, pathologic α-syn binding to TLR2 is involved in the microglia response to pathologic α-syn (*13, 15*).

Although Lag3 is an important protein in the immune system (*35*) and its mRNA is enriched in the brain (*24*) and microglia (*25–27*), we confirmed that Lag3 is not only expressed in microglia, but in neurons through the use of Loxp reporter line with a YFP (yellow fluorescent protein) signal knocked into the Lag3 locus (Lag3^L/L-YFP^) (*28*). Aplp1 mRNA expression is enriched in oligodendrocytes (*36*), while Aplp1 protein has been reported to be restricted to the neuronal surface (*37–40*). Aplp1’s and Lag3’s localization to neurons supports the notion that these two proteins cooperate in the internalization of pathologic α-syn in neurons. Since Lag3 message is enriched in microglia and Aplp1 mRNA is enriched in oligodendrocytes, it will be important to determine the role of Lag3 and Aplp1 in microglia and oligodendrocytes in pathologic α-syn toxicity.

Consistent with our co-immunoprecipitation experiments, NMR analysis confirmed that Aplp1 and Lag3 form a complex where the E1 domain of Aplp1 binds to Lag3 via its D2 and D3 domains. The significantly high chemical shift deviation of the Lag3 D2 domain and Aplp1 E1 domain indicates a direct interaction between Lag3 and Aplp1. α-Syn PFF binds to a common seven amino acid stretch that is contained within the E1, GFLD subdomain of Aplp1 and the D1 domain of Lag3 raising the possibility that there might be a common structural motif that accounts for the binding of pathologic α-syn to Aplp1 and Lag3. Addressing these key questions may facilitate optimization of Aplp1 and Lag3 targeted therapies aimed at pathologic α-syn cell-to-cell transmission.

## MATERIALS AND METHODS

### Study Design

This study aimed to explore the role of Aplp1 and its relationship to Lag3 in the binding, internalization, transmission, and toxicity of pathologic α-syn.

### Animals

C57BL/6 WT were obtained from the Jackson Laboratories (Bar Harbor, ME). *Lag3*^−/−^ mice^40^ were obtained from Dr. Charles G. Drake when he was at Johns Hopkins University. *Aplp1*^−/−^ mice^41^ were obtained from Dr. Ulrike Muller at the University of Heidelberg. Both *Aplp1*^−/−^ and *Lag3*^−/−^ mice were kept in C57BL/6 background (*22, 41–43*). Double knockout of Aplp1 and Lag3 (*Aplp1*^−/−^/*Lag3*^−/−^) were generated by two consecutive crosses. *Aplp1*^−/−^ and *Lag3*^−/−^ were intercrossed to obtain the *Aplp1*^+/–^/*Lag3*^+/–^ mice. These animals were further intercrossed (*Aplp1*^+/–^/*Lag3*^+/–^ × *Aplp1*^+/–^/*Lag3*^+/–^) to obtain double knockout in the next generation (e.g., 6.25% *Aplp1*^+/+^/*Lag3*^+/+^(WT) and 6.25% *Aplp1*^−/−^/*Lag3*^−/−^). All housing, breeding, and procedures were performed according to the guidelines of the NIH and Institutional Animal Care Committee of Johns Hopkins University for the Care and Use of Experimental Animals.

### Generation of α-syn monomer and PFF

Recombinant α-syn protein were purified from ClearColi^TM^ BL21 competent E. coli transformed with full-length α-syn in pRK172 vector. The bacterial endotoxins were removed by Toxineraser endotoxin removal kit (GenScript Biotech Corp., USA) followed with the measurement of the level of endotoxin by ToxinSensor Chromogenic LAL Endotoxin Assay Kit (GenScript Biotech Corp., USA). The recombinant α-syn solution was aliquoted before fibrillization and stored at −80°C until use. Before fibrillization, recombinant α-syn solution was centrifuged at 4°C (12,000 g, 15 min). Supernatant α-syn (5 mg/ml) was transferred into endotoxin-free Eppendorf tubes, and fibrils were generated by shaking for 5-7 days at 37°C with 1,000 RPM Eppendorf Thermomixer. α-Syn PFF were generated by sonication of fibrils with 20% amplitude for a total of 60 pulses (0.5 seconds on/off cycle). The fibrils and PFF were validated by Thioflavin T assay, transmission electron microscopy (TEM), immunoreactivity of anti-pS129 and neurotoxicity in primary neuronal culture.

### Generation of synthetic β-amyloid (1-42) fibrils and PFF

β-amyloid peptide was purchased from Anaspec (AS-23523-05). The β-amyloid peptide was dissolved clearly in Hexafluoro-2-propanol (HFIP), and kept in SpeedVac (Thermo Scientific, USA) for hours until HFIP completely dried down. The β-amyloid film was freshly suspended in DMSO at 2.2 mM and diluted in PBS to obtain a 250 μM stock solution. The β-amyloid fibrils were generated by incubation of the β-amyloid peptide solution in 37°C for 24 hours. The β-amyloid PFF were sonicated with 20% amplitude for a total of 60 pulses (0.5 seconds on/off cycle).

### Transmission electron microscopy (TEM) measurements

Protein samples (∼100 ng/μL) were adsorbed to 400 mesh carbon coated copper grids for 5 min, followed with three times rinse by Tris-HCl (50 mM, pH 7.4). The grids were then floated upon two consecutive drops of 2% uranyl formate, and were dried with filter paper for imaging on a Phillips CM 120 TEM.

### Cell Surface binding assays and ImageJ analysis

SH-SY5Y cells were transfected with related receptors (e.g., Aplp1, Lag3) or mutants. Two days after transfection, the cell cultures with the equivalent cell density were incubated with α-syn-biotin PFF for 1.5-2 hours with different concentrations as indicated in DMEM media with 10% fetus bovine serum (FBS) at room temperature. After incubation, the cells were washed with DMEM media (3 times 5 min), and then fixed with 4% paraformaldehyde (PFA) in PBS for 15 min. After another 3 times wash with PBS, the cells were blocked for 30 min with 10% goat serum and 0.1 % Triton X-100 in PBS. Using alkaline-phosphatase-conjugated streptavidin (1:2000) in PBS supplemented with 5% goat serum and 0.05% Triton X-100, the cells were incubated for overnight (∼16 hours). The bound streptavidin-alkaline phosphatase was visualized by 5-bromo-4-chloro-3-indolyl phosphatase/nitro blue tetrazolium reaction. Bound α-syn-biotin PFF to receptors-transfected SH-SY5Y cells was quantified with ImageJ by normalizing with total cell numbers. Threshold was selected under Image/Adjust in order to achieve a desired range of intensity values for each experiment or image. All the images in each experiment were applied with the same threshold, to exclude the background and obtain the signal. After exclusion of the background, the selected area in the signal intensity range of the threshold was measured using the measurement option under the Analyze/Measure menu. The area values with different α-syn-biotin (monomer or PFF) were input into the Prism program to obtain *K*_d_ or *B*_max_.

### Primary neuronal cultures, α-syn(-biotin) PFF and/or lentivirus transduction, and neuron binding assay

C57BL/6 mice were obtained from the Jackson Laboratories (Bar Harbor, ME). Primary cortical neurons were prepared from E15.5 mouse embryos and cultured in Neurobasal media supplemented with B-27, 0.5 mM L-glutamine, penicillin and streptomycin (Invitrogen, USA) on tissue culture plates coated with poly-L-ornithine. The 3-4 days *in vitro* neuronal cultures were inhibited by F0503-5FU, and were maintained by adding medium every 3 days. Lentiviral cFUGW vectors and particles were generated as previously described (*11, 44*). At 4 days *in vitro*, primary neurons were infected by lentivirus carrying related receptors, or empty vectors as a control [1 × 10^9^ transduction units (TU)/mL] for 72 hours. α-Syn PFF (final concentration 5 μg/mL) were added at 7 days *in vitro* primary neurons, and were incubated for 10–21 days followed by biochemical experiments or toxicity assays. Neurons were harvested for immunofluorescence staining or sequential extraction with Triton X-100 and SDS. Each experiment was performed in duplicate and repeated 3–6 times. To determine the bound signal, α-syn-biotin PFF with different concentrations were administered into primary neuron cultures with the equivalent cell density. Quantification of bound α-syn-biotin PFF to neurons were analyzed with ImageJ by normalizing with total cell numbers.

### Plasmids and deletion mutants

*Lag3* cDNA clones were kindly obtained from Dr. Charles Drake at the Johns Hopkins University, School of Medicine. pcDNA3.1-*Aplp1*, *App* and *Aplp2* cDNA clones were obtained from Dr. Yasushi Shimoda at Nagaoka University of Technology, and Dr. Gopal Thinakaran at the University of Chicago, and Dr. Ulrike Muller at the University of Heidelberg. *Lag3* deletion mutants with a Myc tag, *Aplp1* deletion mutants with a FLAG tag, 7aa substitution mutants of *Lag3*-Myc and Flag-*Aplp1*, and *Aplp1* chimera mutants with a FLAG tag, were constructed by a ligation or In-Fusion method. Briefly, primers were designed to flank the sequences to be deleted or modified. DNA were PCR amplified with herculase polymerase (Agilent Technologies, USA) or CloneAmp HiFi PCR Premix (Clontech, USA). Amplicons were separated on a 1-2% agarose gel and appropriate bands were excised and isolated using a gel extraction kit (Qiagen, USA). These fragments were inserted into the pcDNA3.1-based plasmid or cFUGW lentivirus vector using the T4 ligase (New England Biolabs, USA) or in-Fusion HD cloning kit (Clontech, USA). Vectors were transformed into Stellar competent cells (Clontech, USA) or competent Stbl3 cells (Invitrogen, USA). Integrity of the constructs was verified by sequencing.

Genes encoding human A1E1 (residues 50–146 of *APLP1*), mouse L3D2 (residues 168–256) and L3D3 (residues 257-246) of *Lag3* were synthesized by Union-Biotech (Shanghai) Co, Ltd. Receptor genes were cloned into pET-28a (+), which contains an N-terminal his6-tag for purification.

Full-length human *APLP1* gene with a Flag tag in the N-terminal was cloned from pCAX *APLP1* (Addgene plasmid#30141) and the primers were shown in Key Resources Table (NheI-Flg-*hAPLP*1-F/XhoI-His-*hAPLP1*-R). Based on the NMR results, we substituted nine residues of *APLP1* with alanine: 63L, 64T, 66H, 68D, 84C, 85L, 109Q, 111T, and 114I, and obtained the human FLAG-*APLP1*(mut9) gene with a Flag tag in the N-terminal synthesized by GENEWIZ Bio. Inc. (Suzhou, China). Both of the two genes were ligated into the pcDNA3.1 backbone with *Nhe* I and *Xho* I sites to construct the plasmids pcDNA3.1-h*APLP1* and pcDNA3.1-h*APLP1*(mut9).

Cell membrane localization of Aplp1 and Lag3 mutants: After PBS wash (three times within 5 min), SH-SY5Y cells were incubated for 10 min in 5 μg/mL Wheat Germ Agglutinin, Alexa Fluor™ 488 Conjugate (Invitrogen, W11261) mixed in HBSS. When labeling is complete, the cells were washed in HBSS. For cell membrane localization of Aplp1 and Lag3 mutants, the cells were permeabilized in 0.2% Triton X-100 for 10 mins at room temperature then incubated with anti-FLAG 1:1000 (Cell Signaling Technology, 14793s) and anti-Myc 1:1000 (Cell Signaling Technology, 2276s) for 16 hr at 4°C, then washed and incubated with the secondary antibody.

### Protein Expression, Purification and Refolding

The three constructs encoding A1E1 (residues 50-146), L3D2 (residues 168-256) and L3D3 (residues 257-346) were transformed into *E. coli* BL21 (DE3) cells (Novagen), respectively. The similar protocol for gene expression, protein purification and refolding were used for A1E1, L3D2 and L3D3. Cells were grown to OD_600_ = 1.6, and then induced by 1 mM IPTG. After shaking at 25°C for 12 hr, cells were harvested by centrifugation (5053g, 16 min) and resuspended in a buffer containing 50 mM Tris, 500 mM NaCl at pH 8.0, then lysed by high-pressure homogenizer. The lysate was centrifuged (27, 216 g, 30 min) at 4°C and the pellet was washed in a buffer containing 50 mM Tris, 0.5% Triton X-100 at pH 8.0 and followed by another wash buffer of 50 mM Tris, 1 M NaCl. pH 8.0. Then the inclusion bodies were solubilized in 6 M guanidine hydrochloride containing 50 mM NaCl and 50 mM Tris, pH 8.0. The supernatant was loaded onto a 5-mL Ni-NTA column (GE Healthcare) and eluted with the buffer of 50 mM Na_2_HPO_4_ and 6 M guanidine hydrochloride, pH 4.0. Target protein was further purified by a 300SB-C3 RP-HPLC column (Agilent) with a linear gradient of acetonitrile (0-100%) and then lyophilized.

For the ^15^N-labeled A1E1 protein, the sample preparation protocol was the same as that for unlabeled A1E1 except that the cells over-expressing A1E1 proteins were grown in M9 minimal medium with ^15^NH_4_Cl (1 g/L).

Dry protein powder was solved in a buffer of 50 mM Na_2_HPO_4_, 50 mM NaCl and 6 M Guanidine hydrochloride at pH 7.0 with a maximum concentration of 0.5 mg/mL and filtered by 0.22 um Millipore membranes. Then a three-step dialysis process was performed at 4 °C to fully refold the target protein. Each step takes 3 hours. The buffer used in each step was displayed below. Step 1: 50 mM Na_2_HPO_4_, 50 mM NaCl, 2 M Guanidine hydrochloride, pH 7.0 with 5% glycerin (2 L); Step 2: 50 mM Na_2_HPO_4_, 50 mM NaCl, 1 M Guanidine hydrochloride, pH 7.0 with 2.5% glycerin (4 L); step 3: 50 mM Na_2_HPO_4_, 50 mM NaCl, pH 7.0 (5 L). Then the dialysate was filtered and concentrated before a further purification with a Superdex 75 gel filtration column (GE Healthcare) in a buffer of 25 mM Na_2_HPO_4_, 50 mM NaCl, pH 7.0. Fractions containing the target protein was concentrated and stored at −80 °C. The purity was assessed by SDS-PAGE. Protein concentration was determined by BCA assay (Thermo Fisher).

### NMR spectroscopy

All NMR experiments were collected at 298 K on a Bruker Avance 900 MHz spectrometer equipped with a cryogenic TXI probe. All Protein samples were prepared using the same NMR buffer of 25 mM Na_2_HPO_4_, 50 mM NaCl, and 10% (v/v) D_2_O at pH 7.0. For the titration samples, ^15^N labeled A1E1 was mixed with L3D2/L3D3 at the molar ratio of 1 to 2, and each sample was made to a total volume of 500 μL containing of 35 μM ^15^N-A1E1 in the absence and presence of unlabeled L3D2/L3D3 that diluted from high concentration stocks. Bruker standard pulse sequence (hsqcetfpf3gpsi) was used to collect the 2D ^1^H-^15^N HSQC spectra, with a spectra width of 16 and 30 ppm for proton and nitrogen dimensions. Chemical shift deviations (CSD) were calculated using the equation below,

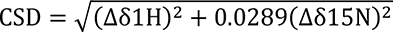

Where Δδ1H and Δδ15N are the chemical shift differences of amide proton and amide nitrogen between free and bound state of A1E1, respectively. All NMR spectra were processed using NMRPipe (*45*) and analyzed using NMRView (*46*).

### Live-cell imaging

α-Syn PFF were labeled with pHrodo dye (Invitrogen, USA), exhibiting minimal fluorescence at neutral pH, and enhanced fluorescence in low pH. WT and Aplp1^−/−^ neurons 9–11 days *in vitro* were infected with lentiviral particles of Aplp1 expression or control, 3 days prior to the addition of α-syn-pHrodo PFF. Live-cell images were recorded by Microscope Axio Observer Z1 (Zeiss, USA) as previously described (*11*). The baseline was established as the fluorescence intensity of the neuron at 2–3 min after α-syn-pHrodo PFF administration. The internalized α-syn-pHrodo PFF at each time point was obtained by tracking individual neuronal objects, images were collected at 30-s intervals. The signal of internalized α-syn-pHrodo PFF was achieved by subtracting the intensity of the baseline in each experiment. All the data acquisition and analysis were performed with the same setting.

### Co-localization of endosome markers and α-syn-biotin PFF

Primary neurons were transfected with FLAG-Aplp1 expression vector 2 days prior to the addition of α-syn-biotin PFF (final concentration 1 μM). After 2-hours incubation, the neurons were fixed and immunostained, and the images were obtained using the same exposure time and treated in the same way for analysis. The signal of α-syn-biotin PFF co-localized with endosome markers (e.g., Rab5, Rab7) was measured and quantified by the Zeiss Zen software as previously described^11^.

### Neuronal internalization of latex beads

Latex beads (Sigma, USA) were applied to 14 days *in vitro* WT, Aplp1^−/−^, Lag3^−/−^, and Aplp1^−/−^/Lag3^−/−^ primary cortical neuron culture (1 μL/mL) for 4-hours incubation at 37°C, as previously described^11^. The numbers of internalized latex beads were quantified by Microscope Axio Observer Z1 (Zeiss, USA).

### Endosome enrichment

α-Syn-biotin PFF were administered to neuron (12–14 days *in vitro*) cultures and incubated for 1.5 hours. Neurons were washed by pre-warmed culture medium followed by 30 sec trypsin to remove the bound α-syn-biotin PFF. Endosomes were enriched as previously described (*11*): the neurons were harvested with PBS and prepared with lysis buffer (250 mM sucrose, 50 mM Tris-HCl [pH 7.4], 5 mM MgCl_2_, 1 mM EDTA, 1 mM EGTA) with a protease inhibitor cocktail (Roche, USA). The suspended cell lysates were pipetted 6 times gently and passed through a syringe 20 times (1 ml TB Syringe, BD, USA), and avoid any bubble in the procedures. The endosomes were harvested in the third pellet followed by three steps of centrifugation 1^st^ (1000 g, 10 min), 2^nd^ (16,000 g, 20 min) 3^rd^ (100,000 g, 1 hr) for immunoblot analysis.

### Uptake of α-syn-biotin PFF in SH-SY5Y cells with Aplp1-Lag3 complex

SH-SY5Y were transfected with pcDNA3.1-h*APLP1* and pcDNA3.1-h*APLP1*(mut9) individually, with/without Lag3 transfection. Two days after transfection, SH cells were incubated with α-syn PFF biotin (31nM or 1 uM) for 2h at 37° C. The cells were treated with 0.25% trypsin for 10 sec after removing the culture medium. The cells were washed three times with PBS, and then fixed with 4% PFA for 15 min. The cells were permeabilized in 0.2% Triton X-100 for 10 min at room temperature following the incubation with anti-FLAG 1:1000 (Cell Signaling Technology, 14793s) and anti-Myc 1:1000 (Cell Signaling Technology, 2276s) for 16 hr at 4° C, then washed and incubated with the secondary antibody.

### Tissue lysate preparation and Western blot analysis

Dissected brain regions of interest or culture samples were homogenized with TX-soluble buffer (50 mM Tris [pH 8.0], 150 mM NaCl, 1% Triton-X 100) containing protease and phosphatase inhibitors (Roche, USA). The supernatants were collected for soluble fraction after centrifugation (20, 000 g, 20 min), and the pellets were resuspended in TX-insoluble buffer (containing 2% SDS) with protease and phosphatase inhibitors. Protein concentrations of samples were determined using the BCA assay (Pierce, USA) and samples (10-20 μg total proteins) were loaded on SDS-polyacrylamide gels (12.5 13.5%) and transferred onto nitrocellulose membranes. Blots were blocked in 5% non-fat milk or 5% BSA in TBS-T (Tris-buffered saline, 0.1% Tween 20) and probed using various primary antibodies and related secondary antibodies, and then were detected using ECL or SuperSignal Femto substrate (ThermoFisher, USA) and imaged by ImageQuant LAS 4000mini scanner (GE Healthcare Life Sciences, USA) or by film.

### Co-immunoprecipitation (co-IP)

For the interaction of Aplp1 and Lag3, brain lysates were prepared, or HEK293FT cells were transfected with receptors of interest or deletion mutants. The cell or brain samples were homogenized with lysis buffer containing 50 mM Tris [pH 8.0], 150 mM NaCl, 1% Triton X-100, and protease inhibitors (Roche, USA). After the centrifugation (20627 g, 20 min), protein concentration of the supernatants was determined using the BCA assay (Pierce, USA). Aliquots of the samples containing 500 µg of protein were incubated with the appropriated microbeads, such as Dynabeads MyOne Streptavidin T1 (Invitrogen, USA), Dynabeads Protein G (Life Technologies, USA) incubated with Myc, 410C9 or CT11 antibody, and FLAG M2 Magnetic Beads, (Sigma, USA). The samples were pre-cleared with the appropriated beads for one hour at room temperature (RT), and meanwhile the related beads were incubated for one hour (10 min for Myc) with the appropriated antibodies and control antibodies. Pre-cleared samples were incubated with microbeads overnight at 4°C or 1hr at RT (for Myc). The IP complexes were washed 4-6 times with IP buffer and then denatured by adding 2x Laemmli Buffer plus β-mercaptoethanol.

### *In situ* Proximity ligation assay (PLA)

Slices were incubated for 1 hr at 37 °C in the blocking solution in a heated humidity chamber, and then with the primary antibodies overnight at 4° C (for mouse Lag3, mouse 410C9, 1:500; For mouse Aplp1, rabbit A1NT, 1:500). The cover slips were washed twice for 5 min with buffer A at room temperature, followed by incubation with the PLA probes (secondary antibodies against two different species bound to two oligonucleotides: anti-mouse MINUS and anti-rabbit PLUS) in antibody diluent in a pre-heated humidity chamber for 60 min at 37° C. Then wash the slides 2 x 5 min in 1x Wash Buffer A at room temperature. In the ligation step, the slides were incubated with the ligation solution in a pre-heated humidity chamber for 30 min at 37° C. The slices were washed with buffer A twice for 5 min before incubation for 100 min with amplification stock solution (contains polymerase) in a heated humidity chamber for 30 min at 37° C. After two washes of 10 min with 1x buffer B and 1min 0.01x buffer B, the slices were mounted with a coverslip using a minimal volume of Duolink® In Situ Mounting Medium with DAPI 15 min before imaging. Fluorescence images were acquired on a Nice confocal laser scanning microscope using a 40X Oil objective for high magnification images.

Image Analysis/quantification of PLA signal: All images analyzed in this experiment were taken from the cortex slices of 2 mice for each genotype. For all experiments, quantifications were performed from 11 images (2 slices for each animal; 2 mice). High-resolution (40X 1.4 NA) images from single scans were analyzed in ImageJ to calculate the density of PLA dots. A same threshold was selected manually to discriminate PLA signals from background fluorescence. The built-in macro ‘Analyze Particles’ was then used to count. Objects larger than 5 μm^2^ were rejected. The remaining signals were counted as PLA signal.

### Microfluidic chambers

Triple compartment microfluidic devices (TCND1000, Xona Microfluidic, LLC, USA) were attached on glass coverslips and coated with 2.5 mg/ml poly-L-lysine overnight and washed with 0.1% Triton X-100 and ddwater three times each, and then replaced with Neurobasal media. Approximately 10,000 neurons were plated per chambers. At 7 days *in vitro*, 0.5 μg α-syn PFF were added in chamber 1 of all the four groups. To prevent the α-syn PFF diffusion due to the flow direction, a 50 μl difference in media volume was maintained between chamber 1 and chamber 2, and chamber 2 and chamber 3 according to the manufacturers’ instructions. Neurons were fixed 14 days after administration of α-syn PFF using 4% PFA in PBS, followed with immunofluorescence staining.

### Cell death assessment

Primary cortical neurons were treated with 5 μg/mL of α-syn PFF for 14–21 days. Percent of cell death was determined by staining with 7 μM Hoechst 33342 and 2 μM propidium iodide (PI) (Invitrogen, USA). PI were diluted in warm neuronal culture medium and incubated for 5 min and then images were taken by a Zeiss microscope equipped with automated computer assisted software (Axiovision 4.6, Carl Zeiss, Dublin, CA). The background signals were subtracted with the intensity of control group, and the percent of neuronal death was calculated by PI signal divided by Hoechst 33342 signal.

### Stereotaxic injection and sample preparation

On the day of intrastriatal injections, α-syn PFF were diluted in sterile PBS and briefly sonicated. Two-three months old mice were anesthetized with a mixture of ketamine (100 mg/kg) and xylazine (10 mg/kg), followed with intrastriatal injection with α-syn PFF (5 μg/2 μL at 0.4 μL/min) at the coordinates of the dorsal striatum: anteroposterior (AP) = +0.2 mm, mediolateral (ML) = +2.0 mm, dorsoventral (DV) = +2.8 mm from bregma. Injections were performed using a 2 μL syringe (Hamilton, USA), and the needle was maintained for an additional 5 min for complete absorption before slow withdrawal of the needle. After surgery, animals were monitored and post-surgical care was provided. Behavioral tests were performed 180 days after α-syn PFF injection, and then mice were euthanized for biochemical and histological studies. For biochemical studies, tissues were immediately dissected and frozen at −80°C. For histological studies, mice were perfused with PBS and 4% PFA and brains were removed, followed by fixation in 4% PFA overnight and transfer to 30% sucrose for cryoprotection.

### Behavioral tests

To evaluate α-syn PFF-induced behavioral deficits, PBS-and α-syn PFF-injected mice were assessed by pole test and cylinder test. The experimenter was blinded to treatment group for all behavioral studies. All tests were recorded and performed between 10:00–16:00 in the lights-on cycle.

### Pole test

Mice were acclimatized in the behavioral procedure room for 30 min. The pole was made of a 75-cm metal rod (diameter 9 mm) and wrapped with bandage gauze. Mice were placed near the top of the pole (7.5 cm from the top of the pole) facing upwards. The total time taken to reach the base of the pole was recorded. Before the actual test, mice were trained for three consecutive days. Each training session consisted of three test trials. On the test day, mice were evaluated in three sessions and the total time was recorded. The maximum cutoff time to stop the test and recording was 120 sec. Results for turn down, climb down and total time were recorded.

### Cylinder test

Spontaneous movement was measured by placing animals in a small transparent cylinder (height, 15.5 cm; diameter, 12.7 cm). Spontaneous activity was recorded for 10 min. The number of forepaw touches, rears and grooming were measured. Recorded files were viewed and rated in slow motion by an experimenter blinded to the mouse type.

### Purification of anti-Lag3 410C9 and labeling

410C9 hybridoma cell line is available from Dr. Dario Vignali under a material agreement with the University of Pittsburgh. We cultured the 410C9 hybridoma cell line, and collected the supernatant culture medium. The supernatant was concentrated and purified over HiTrap Protein G HP antibody purification column (GE Healthcare). To visualize 410C9 in vivo, a near infrared (IR) dye IRDye680RD was used to label 410C9. The 410C9 solution was mixed with IRDye680RD NHS Ester (molar ratio 1:2). The mixture solution was consistently shaking on the shaker at 4°C for 16 hrs, and then dialyzed against DI water for 24 hrs using the dialysis membrane with molecular weight cut-off of 3,500 Da to remove any free dye. The obtained solution was subjected to BCA assay to determine the concentration of 410C9 for further study.

### Quantification and Statistical Analysis

All data were analyzed using GraphPad Prism 8. Statistics Data are presented as the mean ± SEM or mean ± SD. with at least 3 independent experiments. Representative morphological images were obtained from at least 3 experiments with similar results. Statistical significance was assessed via a one or two-way ANOVA test followed by indicated post-hoc multiple comparison analysis. Assessments with *p* < 0.05 were considered significant.

## Acknowledgements

We thank I.-H. Wu for graphic art assistance. We appreciate the Aplp1 antibodies as gift provided by Dr. Gopal Thinakaran at the University of South Florida Morsani College of Medicine and the support from Dr. Haiquan Mao at the Johns Hopkins University. The authors acknowledge the join participation by the Adrienne Helis Malvin Medical Research Foundation through its direct engagement in the continuous active conduct of medical research in conjugation with the Johns Hopkins Hospital and the Johns Hopkins University School of Medicine and the Foundation’s Parkinson’s Disease Program M-2014. T.M.D. is the Leonard and Madlyn Abramson Professor in Neurodegenerative Diseases.

## Funding

National Institutes of Health Grant R01 NS107318 (XM)

National Institutes of Health Grant R01NS107404 (HSK)

National Institutes of Health Grant K01 AG056841 (XM)

National Institutes of Health Grant P01AI108545 (DAAV, CJW)

National Institutes of Health Grant R01 AI144422 (DAAV, CJW)

Parkinson’s Foundation the Stanley Fahn Junior Faculty Award PF-JFA-1933 (XM)

Maryland Stem Cell Research Foundation Discovery Award 2019-MSCRFD-4292 (XM)

American Parkinson’s Disease Association 90076052 (XM)

Uehara Memorial Foundation, Japan (YK)

JPB Foundation (TMD)

Adrienne Helis Malvin Medical Research Foundation, M2014 (TMD, VLD)

Parkinson’s Disease Foundation, PDF-SFW-1572 (PG)

American Parkinson Disease Foundation, PDF-APDA-SFW-1650 (PG).

## Author contributions

XM, HG, DK, led the project and contributed to all aspects of the study. YK, EX, HW, CC, YL, HB, LJ, NW, MC, AL, JY, CR, MS, PG, SB, SK, Shu Z, contributed to biochemical, cellular and mouse experiments. SS. contributed to HPLC analysis. XK, YS, HM, CJW, DAA, UCM provided key reagents. Shengnan Z, CJ, and CL contributed to NMR studies. XB, HSK, VLD, and TMD designed research, XM, HG, DK, YK, EX, HSK, VLD, and TMD. analyzed data, and XM, VLD, and TMD. wrote the paper. All authors reviewed, edited and approved the paper.

## Data and materials availability

Further information and requests for resources and reagents should be directed to and will be fulfilled by Ted M. Dawson. There are no restrictions on any data or materials presented in this paper. All data are available in the main text or the supplementary materials.

## Supplementary Materials

**Fig. S1.**
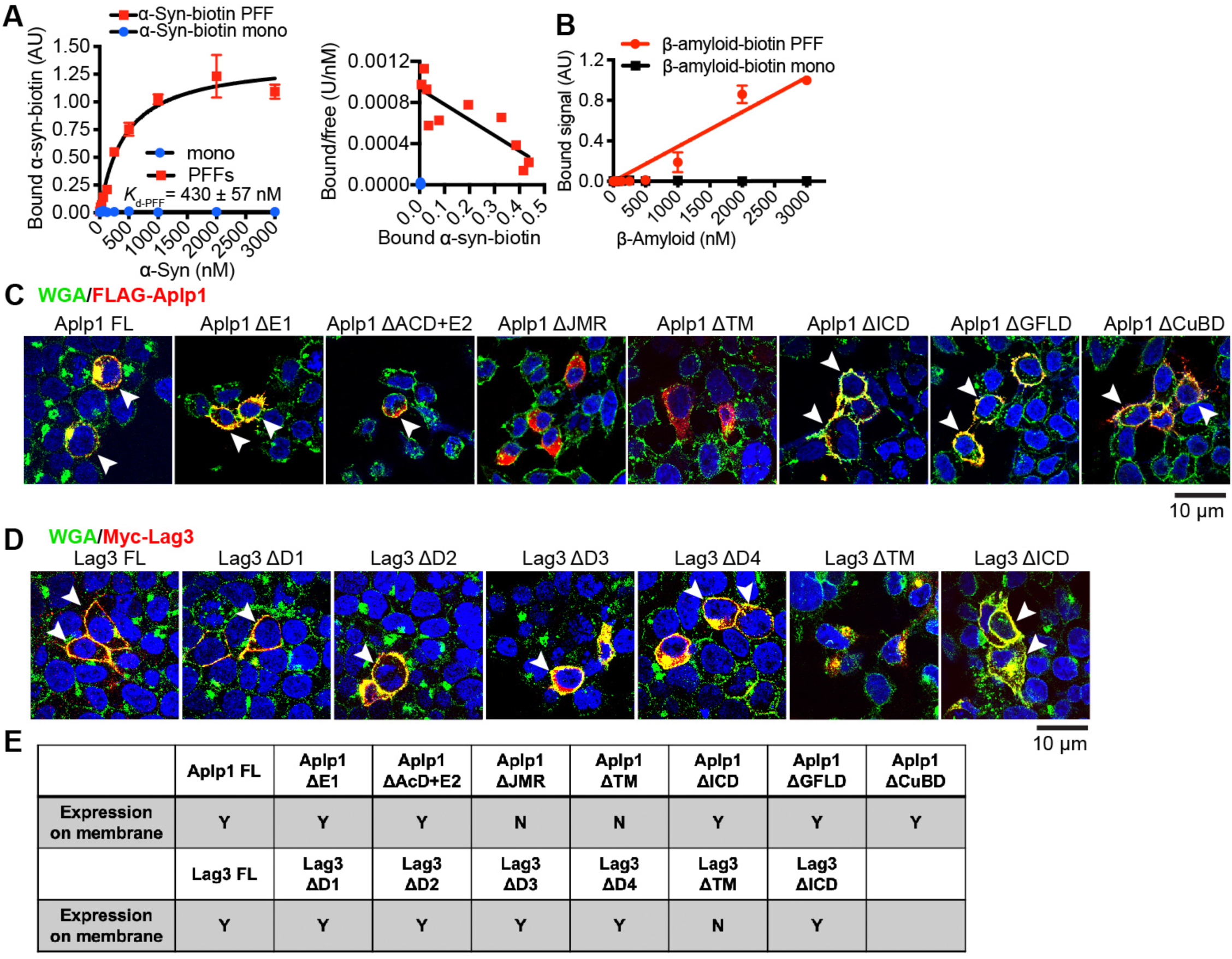
Aplp1 binds to α-syn-biotin PFF but not monomer. **(A)** α-Syn-biotin PFF binding to SH-SY5Y cells expressing Aplp1 as a function of α-syn concentration with Scatchard analysis. α-Syn-biotin monomer has minimal binding affinity with cells. *K*_d-PFF_ = 430 ± 57 nM for PFF, data are the means ± SEM, *n* = 3 independent experiments. **(B)** β-Amyloid PFF binds to Aplp1 in non-specific manner (*K*_d_ > 3000 nM), but β-amyloid monomer does not exhibit appreciable binding to Aplp1. **(C, D)** The membrane localization of Aplp1 mutants and Lag3 mutants. **(C)** The anti-FLAG immunostaining images of Aplp1 FL, ΔE1, ΔAcD+E2, ΔJMR, ΔTM, ΔICD, ΔGFLD, ΔCuBD with WGA-488. White arrows indicate the co-localization of FLAG-Aplp1 and WGA. **(D)** The anti-Myc immunostaining images of Lag3 FL, ΔD1, ΔD2, ΔD3, ΔD4, ΔTM, ΔICD, using membrane marker wheat germ agglutinin (WGA) conjugated with Alexa Fluor^TM^ 488. White arrows indicate the co-localization of Myc-Lag3 and WGA. **(E)** The detailed information for the expression on membrane.

**Fig. S2.**
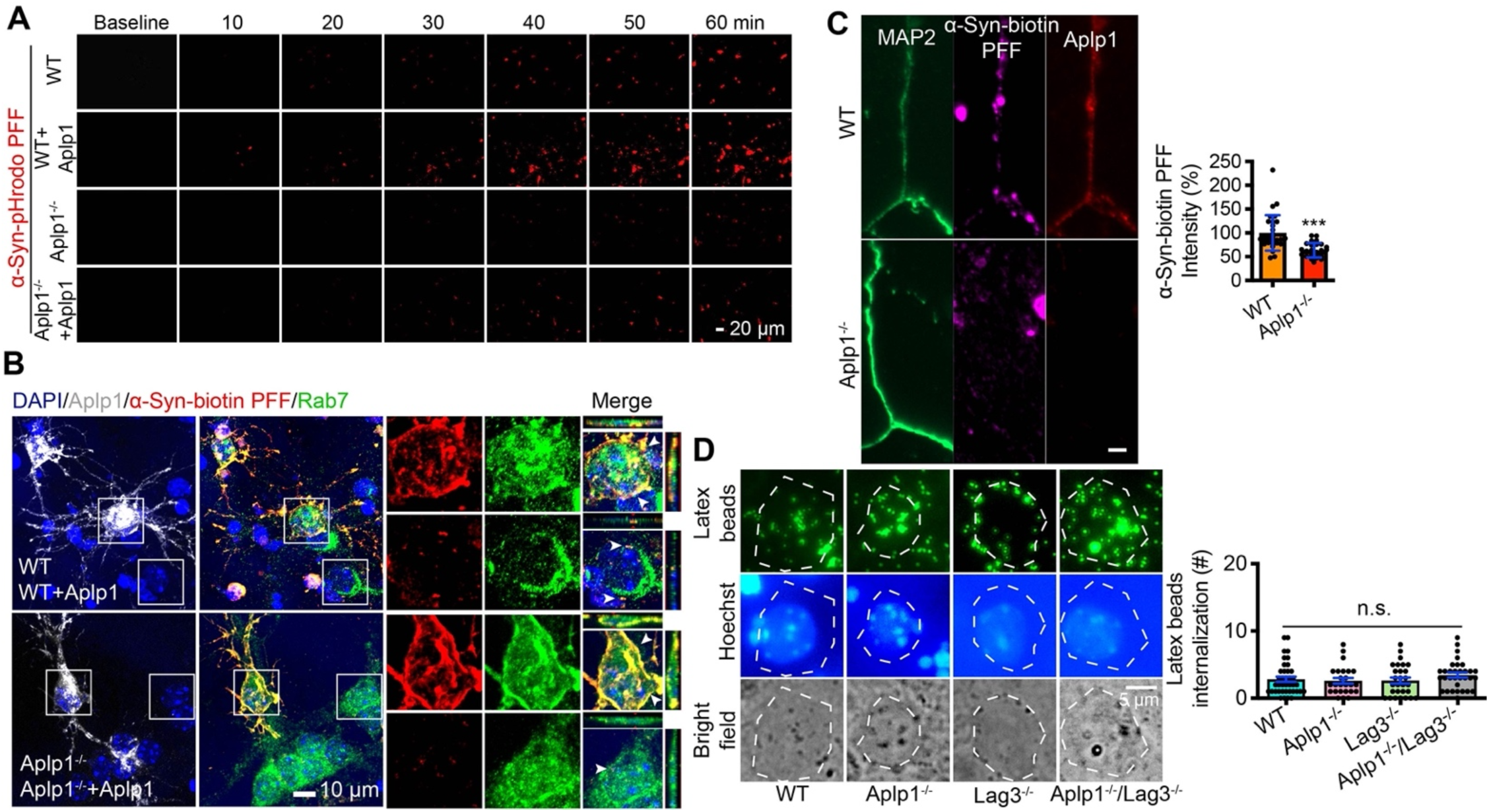
Aplp1 mediates the endocytosis of α-syn PFF. **(A)** Live images of the endocytosis of α-syn-pHrodo PFF. α-Syn PFF were conjugated with a pH-dependent dye (pHrodo red), in which fluorescence increases as pH decreases from neutral to acidic environments. Four groups include wildtype (WT) neurons, WT neurons overexpressing Aplp1 by lentivirus transduction (WT+Aplp1), *Aplp1^−/−^* neurons, and Aplp1^−/−^ neurons with Aplp1 overexpression (*Aplp1^−/−^*+Aplp1). Scale bar, 20 μm. **(B)** The co-localization of internalized α-syn-biotin PFF (red), Rab7 (green) and FLAG-Aplp1 (grey scale) in soma of WT and Aplp1^−/−^ neuronal culture was assessed by means of confocal microscopy. Scale bar, 10 μm. **(C)** Co-localization analysis of α-syn-biotin PFF (red) with MAP2 (green), WT (30 neurites) and Aplp1^−/−^ (28 neurites) from three independent experiments. Data are as means ± SD. Statistical significance was determined by using Student’s *t*-test; \*\*\**P* < 0.001. **(D)** Latex bead internalization in WT, *Aplp1*^−/−^, *Lag3*^−/−^ and *Aplp1^−/−^*/*Lag3*^−/−^ neurons. WT (42 cells), *Aplp1*^−/−^ (32 cells), *Lag3*^−/−^ (32 cells), *Aplp1*^−/−^/*Lag3*^−/−^ (32 cells) from *n* = 3 individual experiments. Scale bar, 5 μm. Quantification of the latex bead internalization. Data are as means ± SD. n.s., not significant. One-way ANOVA followed by Tukey’s correction.

**Fig. S3.**
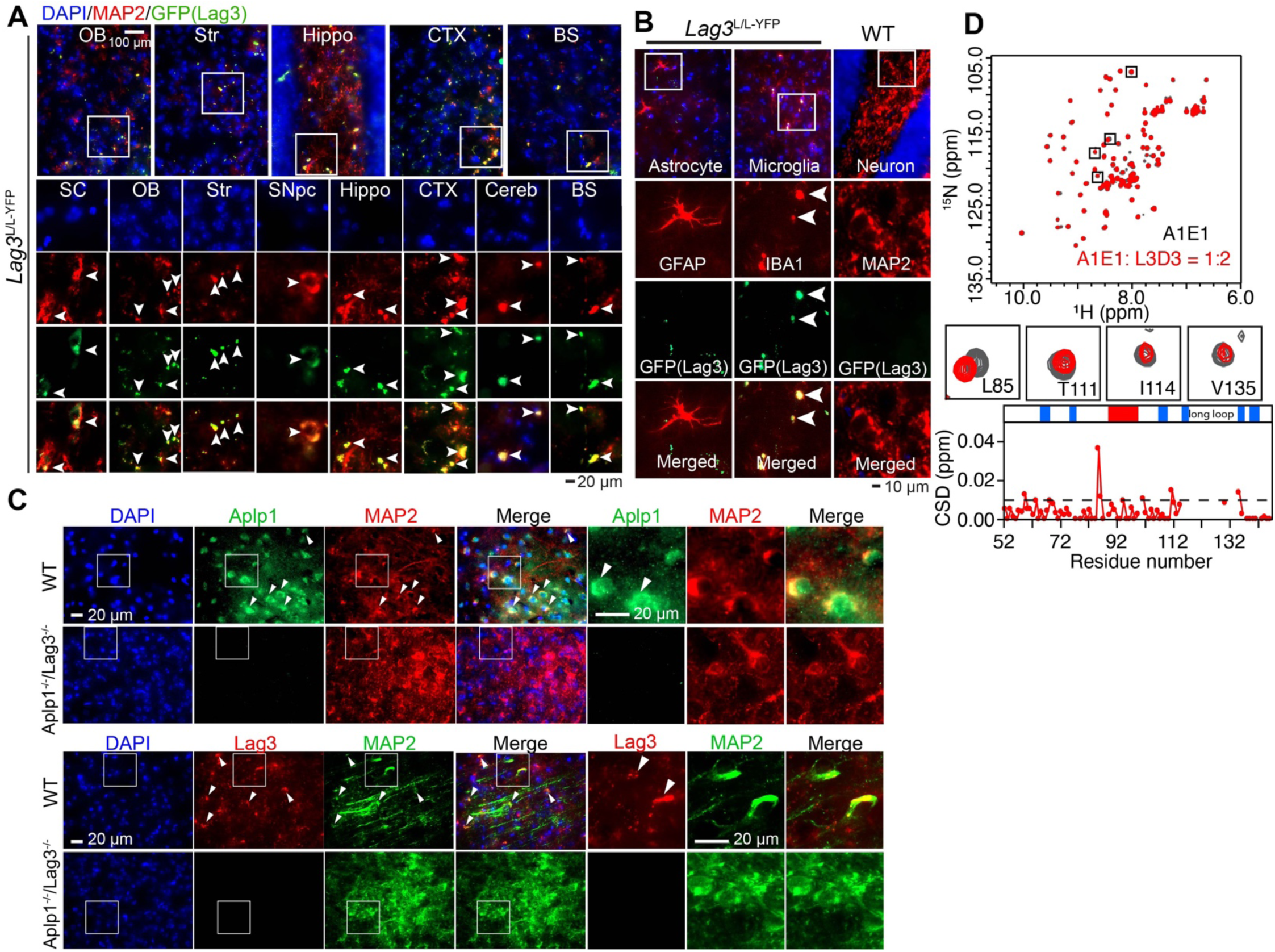
Lag3 is expressed in neurons. **(A)** Cellular localization of Lag3 in a Lag3 Loxp reporter line with a YFP (yellow fluorescence protein) signal knocked into the Lag3 locus (*Lag3*^L/L-YFP^). The immunoreactivity of anti-GFP (Lag3), which recognizes YFP co-localized with MAP2 -positive cells (neurons) in the spinal cord (SC), olfactory bulb (OB), striatum (Str), substantia nigra pars compacta (SNpc), hippocampus (Hippo), cortex (CTX), cerebellum (Cereb) and brainstem (BS). The white arrows indicate the co-localization of GFP(Lag3) and MAP2. Scale bar, 100 μm. **(B)** Co-localization of anti-GFP (Lag3) signals with IBA1-postivie cells (microglia), but not with GFAP-positive cells (astrocyte). The white arrows indicate the co-localization of GFP(Lag3) and IBA1, but not GFAP. Scale bar, 10 μm. **(C)** Immunostaining of Aplp1 and Lag3 in the cortex region of WT and *Aplp1*^−/−^ /*Lag3*^−/−^ mice, with anti-MAP2 immunostaining. **(D)** A1E1 interacts with L3D3. (Upper inset): Overlay of the 2D ^1^H-^15^N HSQC spectra of A1E1 alone (black) and in the presence of 2 molar folds of L3D3 (red). (Middle inset): The same 4 residues that with significant CSDs (> 0.03 ppm) upon L3D2 titration are highlighted in the black boxes and zoomed in. Histogram of the CSDs of A1E1 in the presence of L3D3 at a molar ratio (A1E1/L3D3) of 1:2. (Bottom inset): The domain organization of A1E1 is indicated on the top, with blue boxes indicating the *β*-strands and the red box indicating the *α*-helix. A dashed line was drawn to highlight the residues with CSDs > 0.01 ppm.

**Fig. S4.**
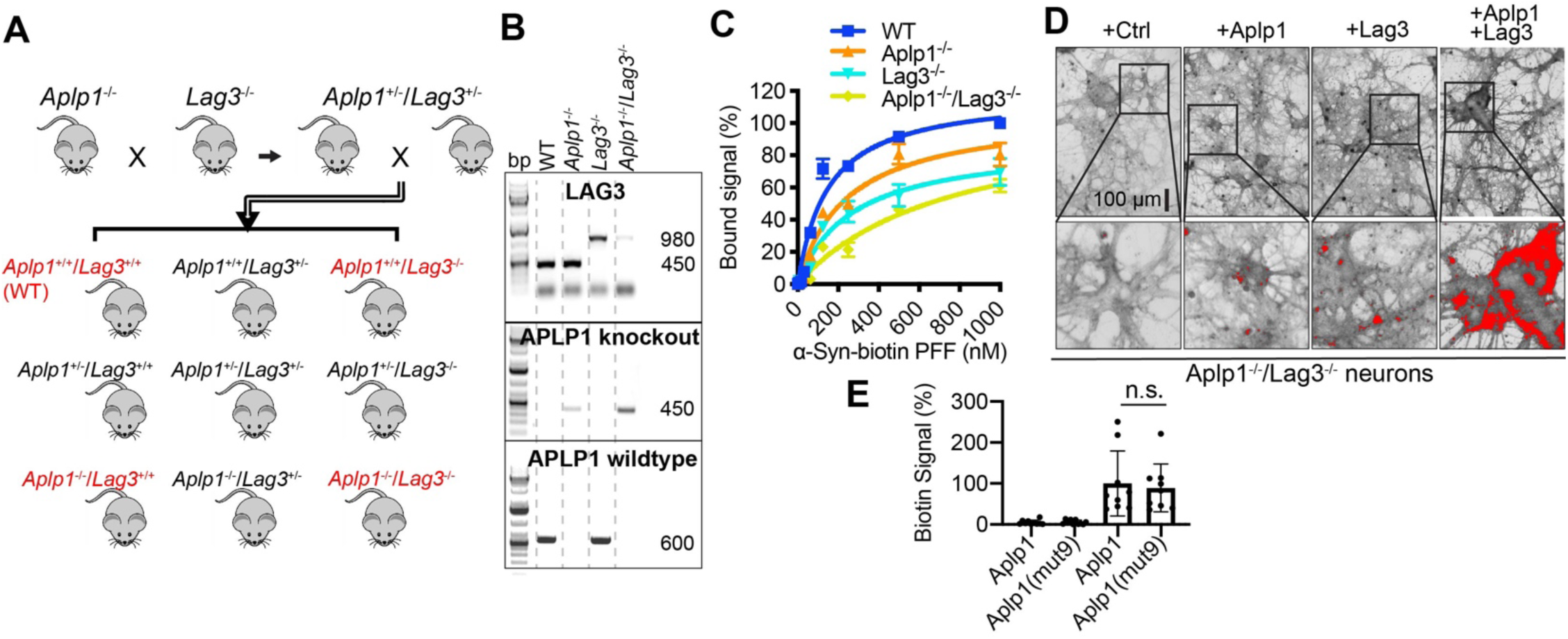
The Aplp1-Lag3 complex and α-syn PFF binding. **(A)** Breeding strategy of the double knockout of *Aplp*1 and *Lag3* (*Aplp1*^−/−^/*Lag3*^−/−^) mice. **(B)** Genotyping confirmation of WT, *Aplp1*^−/−^, *Lag3*^−/−^ and *Aplp1*^−/−^/*Lag3*^−/−^ mice. **(C)** Binding signal of α-syn-biotin PFF to WT, *Aplp1*^−/−^, *Lag3*^−/−^, and *Aplp1*^−/−^/*Lag3*^−/−^ cortical neurons, normalized by the total cell numbers. WT-*K*_d_=149 nM, *Aplp1*^−/−^-*K*_d_=245 nM, *Lag3*^−/−^-*K*_d_=266 nM, *Aplp1*^−/−^/*Lag3*^−/−^-*K*_d_=767 nM. Data are the means ± SEM, *n* = 3 independent experiments. **(D)** Binding images of α-syn-biotin PFF (63 nM) to *Aplp1*^−/−^/*Lag3*^−/−^ neurons transduced with Aplp1, Lag3, or Aplp1+Lag3 by lentivirus, normalized by the total cell numbers. The binding intensity (red signal) was extracted with the ImageJ: Image/Adjust/Threshold setting value 80. Scale bar, 100 μm. **(E)** The uptake signal of α-syn-biotin PFF (31 nM and 1 μM) in Aplp1 and Aplp1(mut9) transfected cells.

**Fig. S5.**
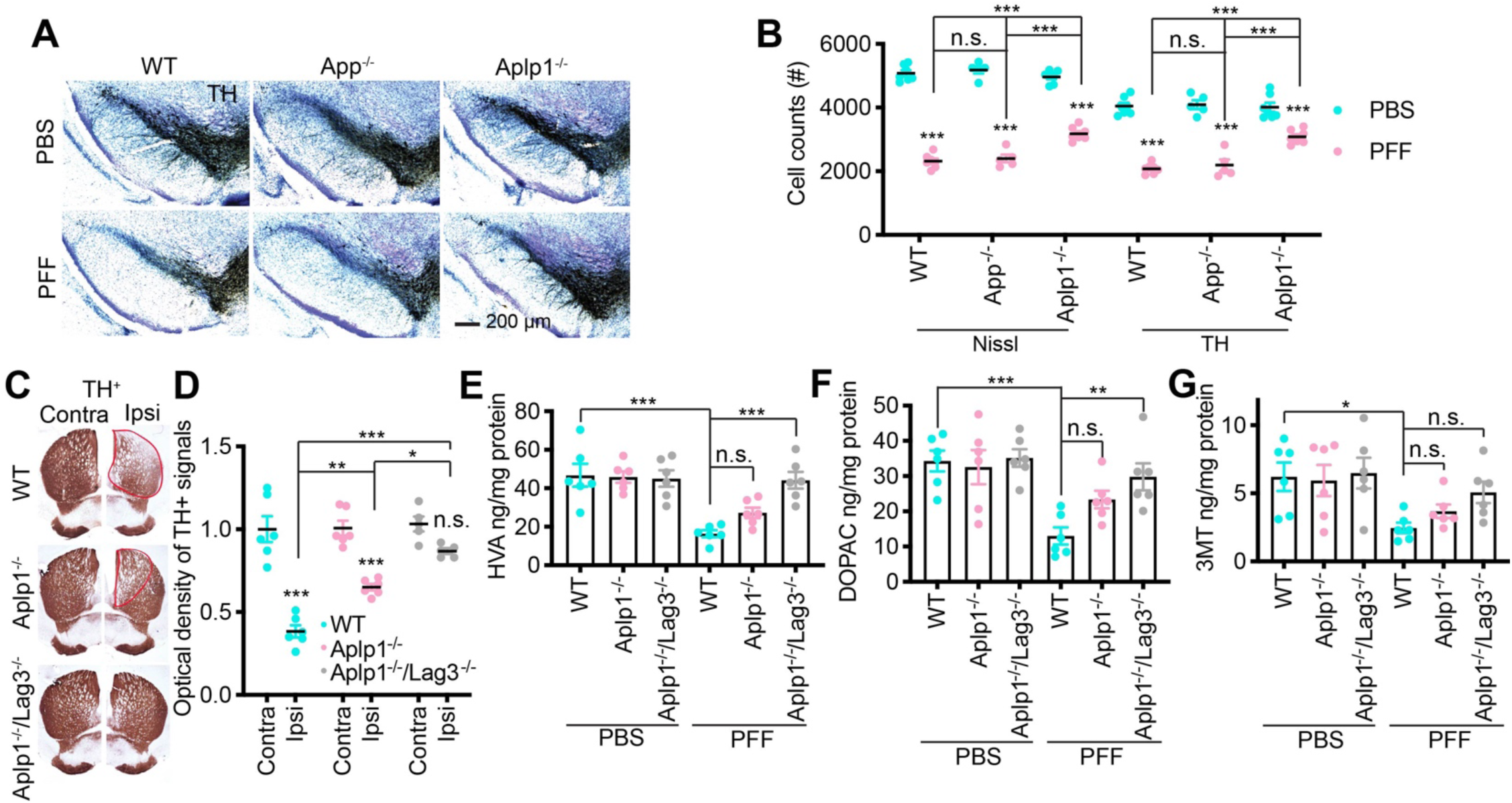
Deletion of Aplp1 and the Aplp1-Lag3 complex prevent neurodegeneration induced by α-syn PFF, but not APP deletion. **(A)** Representative TH and Nissl staining images in the SNpc of α-syn PFF-injected hemisphere in the WT, *App*^−/−^, and *Aplp1*^−/−^ (WT: *n* = 7; *App*^−/−^: *n* = 5; *Aplp1*^−/−^: *n* = 7). **(B)** Stereological counting of the number of Nissl- and TH-positive neurons in the substantia nigra via unbiased stereological analysis after 6 months of α-syn PFF injection in the WT, *App*^−/−^, and *Aplp1*^−/−^ mice. Data are the means ± SEM. Statistical significance was determined by using one-way ANOVA followed with Tukey’s correction; \*\*\**P* < 0.001, n.s., not significant. **(C)** Representative TH immunohistochemistry images in the striatum of α-syn PFF injected brain of WT, *Aplp1*^−/−^, and *Aplp1*^−/−^/*Lag3*^−/−^ mice. **(D)** Quantifications of TH-immunopositive fiber densities in the striatum (WT: *n* = 6; *Aplp1*^−/−^: *n* = 6; *Aplp1*^−/−^/*Lag3*^−/−^: *n* = 5). Data are the means ± SEM. Statistical significance was determined by using one-way ANOVA followed with Tukey’s correction, \**P* < 0.05, \*\**P* < 0.01, \*\*\**P* < 0.001, n.s., not significant. **(E–G)** Striatal metabolites levels in WT, *Aplp1*^−/−^, and *Aplp1*^−/−^/*Lag3*^−/−^ mice, were measured at 180 days post-injection by HPLC. Data are the means ± SEM, *n* = 6 mice per group, one-way ANOVA with Dunnett’s correction.

**Fig. S6.**
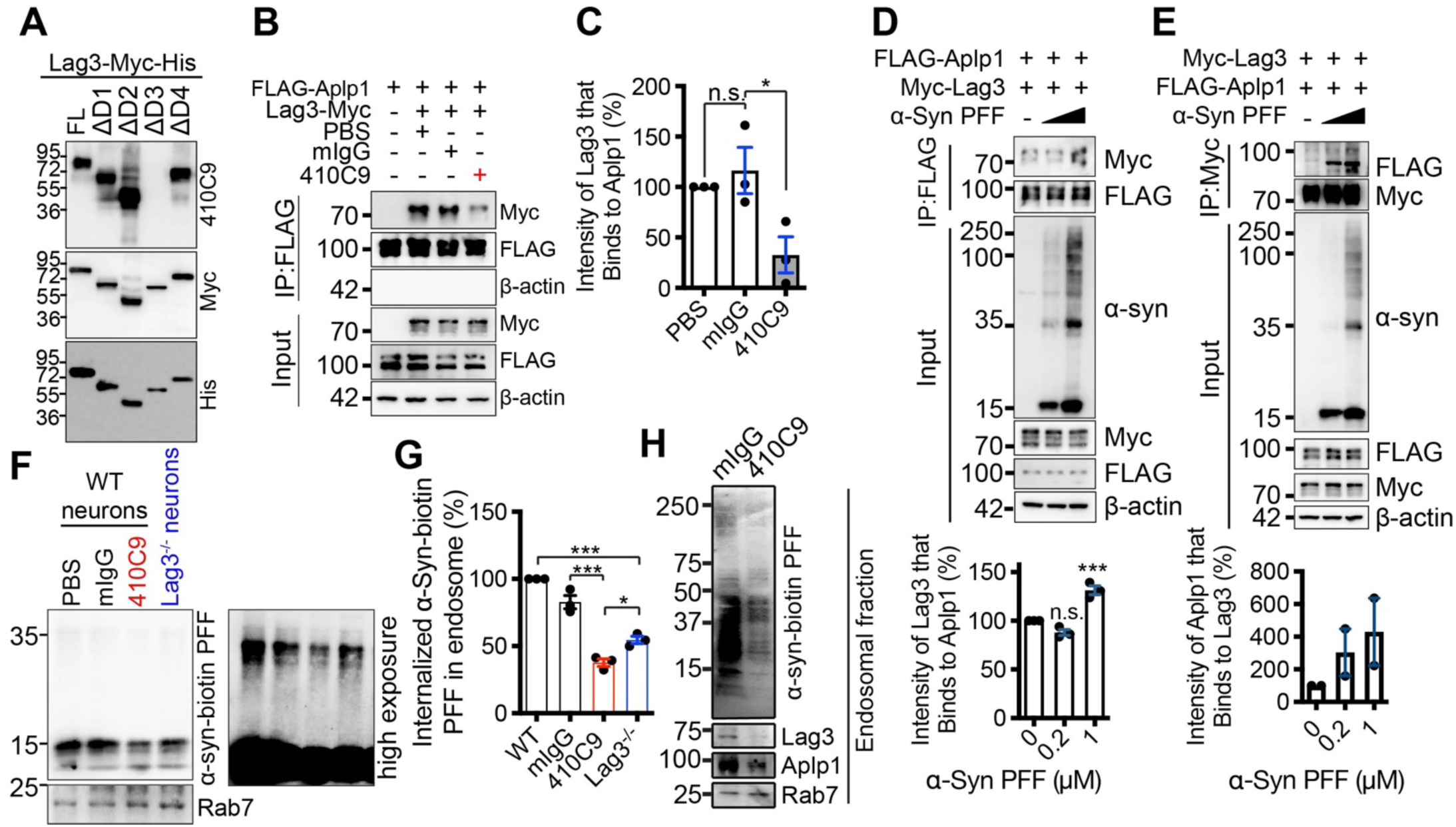
Anti-Lag3 410C9 disrupts the interaction of LAG3 and Aplp1 and LAG3 and Aplp1 co-bind α-syn PFF. **(A)** Epitope of anti-Lag3 410C9 for mouse Lag3. Deletion mutants of Lag3 with Myc and His tags, were transfected in HEK293FT cells, and the cell lysis was probed with 410C9, anti-Myc and anti-His antibodies. Anti-Lag3 410C9 can distinguish the D3 domain of mouse Lag3. **(B and C)** Anti-Lag3 410C9 significantly disrupts the co-IP of FLAG-Aplp1 and Myc-Lag3 in HEK293FT cells. One-way ANOVA followed by Tukey’s correction. **(D**) α-Syn PFF treatment of the cellular extract of FLAG-Aplp1 and Myc-Lag3 transfected HEK293FT cells significantly increases the co-IP of FLAG-Aplp1 and Myc-Lag3. Data are the means ± SEM, \*\*\**P* < 0.001, n.s., not significant; one-way ANOVA followed by Tukey’s correction. *n* = 3 independent experiments. **(E)** α-Syn PFF increases the co-IP of Myc-Lag3 and FLAG-Aplp1. *n* = 2 independent experiments. **(F and G)** 410C9 blocks the internalization of α-syn-biotin PFF significantly more in WT cultures than in *Lag3*^−/−^ cultures. Quantification of the intensity of internalized α-syn-biotin PFF normalized by Rab7. One-way ANOVA followed by Tukey’s correction. **(H)** Both the levels of the endosomal Aplp1 and Lag3 were decreased by 410C9 in WT cortical neuron cultures. Data are the means ± SEM, \**P* < 0.05, \*\**P* < 0.01, \*\*\**P* < 0.001; n.s., not significant.

**Fig. S7.**
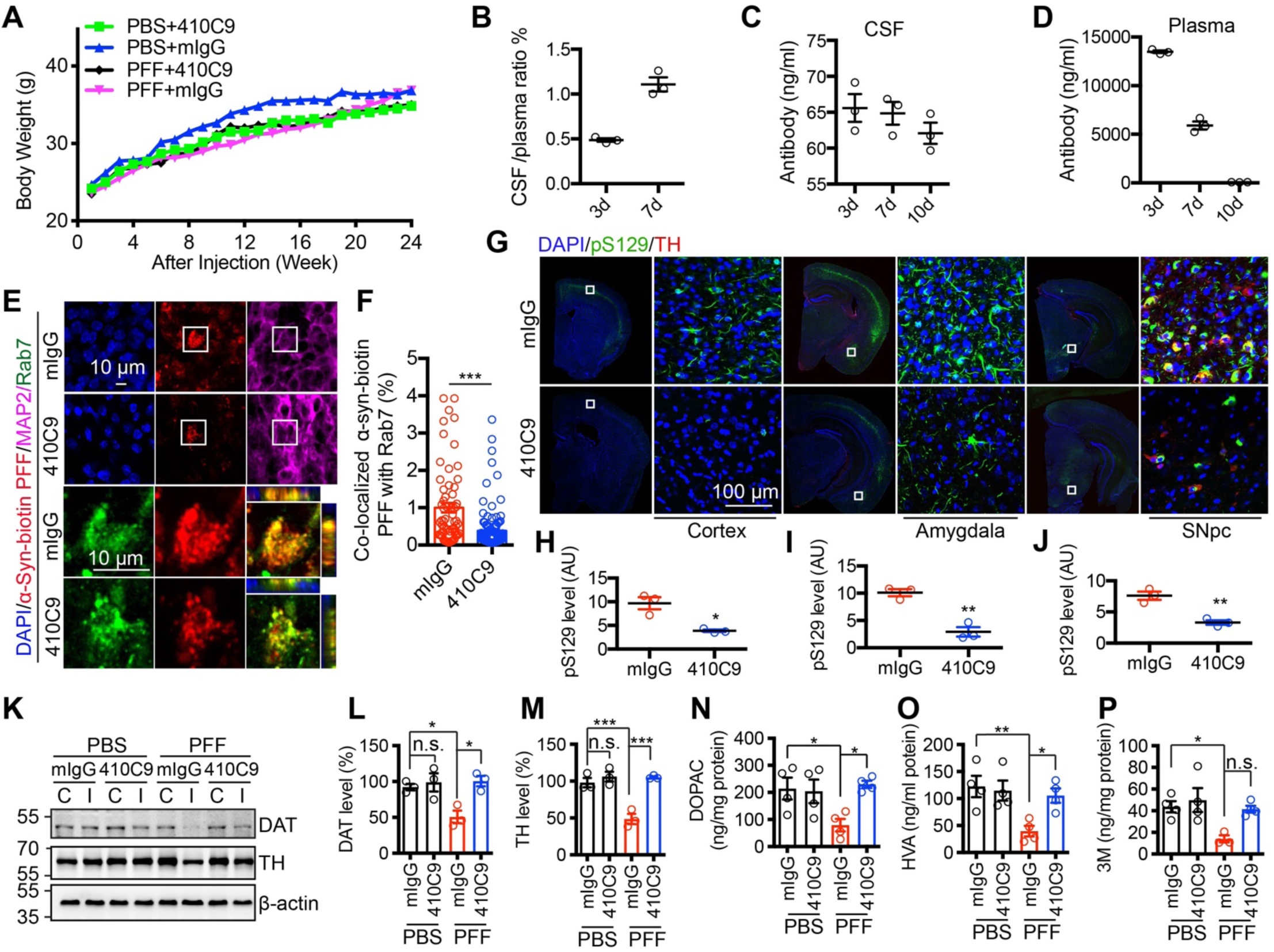
Anti-Lag3 410C9 blocks neurodegeneration induced by α-syn PFF *in vivo*. **(A)** Body weight assessment of WT mice for 6-months after stereotaxic injection of α-syn PFF or PBS, and i.p.(intraperitoneal) treated with 410C9 or mIgG. **(B)** Ratio of 410C9 detected in CSF vs plasma was roughly 0.5% at 3 days. *n* = 3 mice per each time point. Values are means ± SEM. **(C and D)** 410C9 turnover in WT mice. At 3 days, 7 days, and 10 days following a single i.p. injection of 410C9 (10 mg/kg), WT mice (without α-syn PFF treatment) were sacrificed and their (**C**) CSF and (**D**) plasma were collected to determine the level of circulating 410C9. The concentration of near infrared dye (IR) labeled 410C9 was measured by plate reader. **(E and F)** Co-localization of α-syn-biotin PFF with Rab7 is inhibited by 410C9 *in vivo*. The signal of α-syn-biotin PFF colocalizes with Rab7 in neurons (MAP2 staining) in the striatum of WT mice. Intrastriatal injection of α-syn-biotin PFF were performed 3 days after i.p. injection of mIgG or 410C9 (10 mg/kg) in WT mice. Scale bar 10 μm. Data are the means ± SEM, *n* = 3 mice per group, mIgG (63 cells) and 410C9 (112 cells). Statistical significance was determined by using Student’s *t*-test; \*\*\**P* < 0.001. **(G–J)** The representative images of pS129 immunostaining from α-syn PFF-injected WT mice with treatment of 410C9 or mIgG. α-Syn PFF-induced pathology is reduced by 410C9. Quantification of pS129 intensity in (H) cortex, (I) amygdala, and (J) SNpc region. Bars represent mean ± SEM, *n* = 3 mice per group. Statistical significance was determined by using Student’s *t*-test; \**P* < 0.05, \*\**P* < 0.01. **(K, L, and M)** Immunoblot and quantification analysis of the striatum from α-syn PFF or PBS in WT mice, using antibodies against DAT (dopamine transporter) and TH. Mean intensity values are the ratios of the ipsilateral (I) injected side to contralateral (C) side for DAT and TH. Data are the means ± SEM, *n* = 3 mice per group, one-way ANOVA with Dunnett’s correction. \**P* < 0.05, \*\*\**P* < 0.001, n.s., not significant. **(N, O, and P)** Striatal metabolites levels of DOPAC, HVA, and 3M were measured by HPLC-ECD. Data are the means ± SEM, *n* = 4 mice per group, one-way ANOVA with Tukey’s correction; \**P* < 0.05, \*\**P* < 0.01, n.s., not significant.

